# Relating strain amplitude, strain threshold and bone formation rate to exogenous forcing frequency

**DOI:** 10.1101/2023.10.08.561406

**Authors:** Jitendra Prasad, Achsah Marlene Aruva, Himanshu Shekhar, Sanjay Singh, Satwinder Jit Singh

## Abstract

The literature supports the existence of a strain threshold, above which cortical bone adapts to exogenous mechanical loading by forming new bone. This strain threshold, however, varies with loading conditions, locations, waveforms, frequency etc. and there is a need to mathematically express the strain threshold in terms of these parameters. There have been several parametric, mathematical or numerical models in the literature for the cortical bone’s adaptation to mechanical loading, which may be already fitting some of the experimental data; however, they may not be easily and confidently derived from the first principles. To fill the gap, this work has attempted to derive the corresponding bone formation rate (BFR) rather from the first principles, namely using the energy principles. The derived model has been compared to the existing parametric models and validated with respect to the diverse experimental data available in the literature. The developed model is able to not only predict the BFR, but also helps to understand the nature and possible mathematical form of the strain threshold for cortical bone’s adaptation to mechanical loading.

## 1. Introduction

It has been long established through Wolff’s law, that the bone adapts to exogenous mechanical loading [1]. Frost’s mechanostat theory proposed a strain threshold above which there is a new bone formation for the bone to adapt to an exogenous mechanical loading [2]. Strain threshold has been supported by many works in the literature [3]. This strain threshold was, however, found to vary with loading conditions, locations, loading waveform, frequency of loading etc. [4], [5], [6], [7], [8], [9], [10]. For example, the strain thresholds reported for the cantilever loading of mouse tibia, with trapezoidal loading waveform are about 650 με [4] and 850 με [5], with and without rest-inserted waveforms, respectively. The strain threshold in case of four-point loading of rat tibia was found to be higher, such as 1050 με [6]. In case of ulnar axial loading, the strain threshold was found to vary with frequency, from about 1820 με to approximately 650 με as the frequency of the loading waveform increased from 1 Hz to 10 Hz [7]. For such loading, the threshold was also found to vary approximately between 1343 με and 3074 με depending on the location [8]. For axial loading of tibia, the strain threshold was found to be approximately 1100 με [9]. Bone formation rate (BFR) was further found to depend on axial preloads, with higher BFR for lower axial preload, possibly due to preload-dependent strain threshold [10].

Despite its complex nature, strain threshold is an important parameter to quickly ascertain if some physical intervention will induce new bone formation at all. Moreover, some bone adaptation models (e.g. [11], [12]) directly use strain thresholds to compute site-specific mineral apposition rate (MAR) or an average bone formation rate (BFR) at a bone cross-section. Accuracy of strain threshold will directly enhance accuracy of such adaptation models. While there has been strong evidence of strain threshold’s dependance on a number of parameters such as loading condition, location, loading waveform, frequency etc., there is no existing mathematical model in the literature that clearly and elegantly relates the strain threshold to these parameters. The present work attempts to fill the gap by deriving the strain threshold from first principles and validating with the experimental data.

The variable nature of the strain threshold may indicate that the mechanical strain may not be directly inducing new bone formation. There have been accordingly diverse mathematical or computer models of cortical bone’s adaptation to mechanical loading. For example, thresholds for new bone formations have been popularly suggested in terms of strain-energy density [13], [14], [15]. Threshold of viscous dissipation energy can also be found in the literature for modeling bone adaptation [16]. Other kinds of thresholds in the literature include principal stress [13], fatigue damage accumulation [12], [13], strain rate (i.e. strain times frequency, [7], [17]), strain gradient [18], etc. As such, there is no consensus on the nature of threshold for loading-induced new bone formation. Derivation of threshold from first principles may help to identify the correct nature of the threshold and the current work is a step towards such identification.

The present work first proposes an equivalent thermodynamic model to derive how bone formation rate (BFR) can possibly be related to viscoelastic energy dissipated during loading. The derived model has been compared to existing models of bone adaptation and has been successfully validated using the experimental data in the literature (e.g. [7], [19]). The model brings out a mathematical expression of strain threshold as a direct or indirect function of loading, location, loading waveform, frequency, preload etc. The obtained expression for strain threshold varies with the parameters, in accordance with the experimental data found in the literature.

The rest of the paper is organized as follows. Derivations of BFR and strain threshold are given in Section 2, i.e. the Methodology section. Related computational methodologies are also given in Section 2. The model has been validated and the results have been presented in Section 3. The results have been discussed in Section 4. The last section, i.e. Section 5 summarizes the findings and importance of this work.

## 2. Methodology

### 2.1. The 1D Poroelastic Model

Biot [20] derived the settlement of the poroelastic column of height *L* subjected to a compressive stress of *σ*_0_ as a function of time (*t*) as follows:

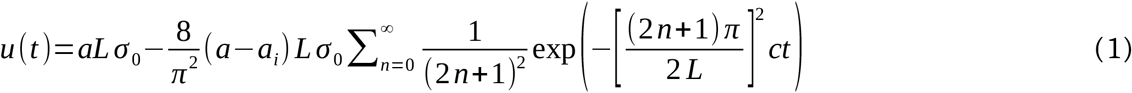

where *a* = final compressibility, *a*_*i*_ = instantaneous compressibility and *c* = consolidation constant are properties of the poroelastic material as defined by Biot [20].

The corresponding creep is given by

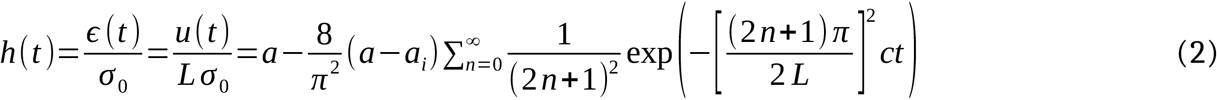

Taking Laplace Transform of the above:

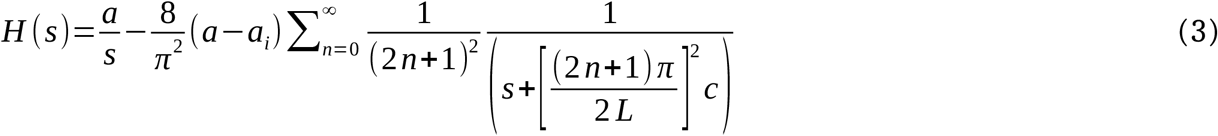

where *s* is the complex frequency.

The corresponding modulus in complex frequency domain is

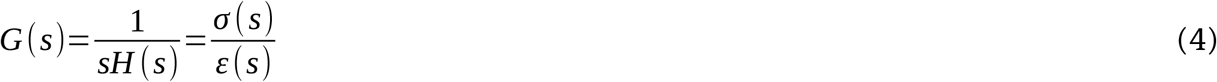

or

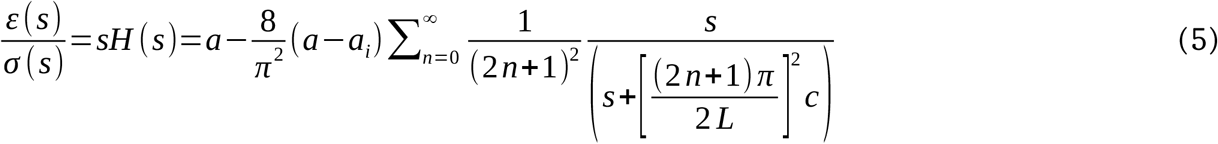

or

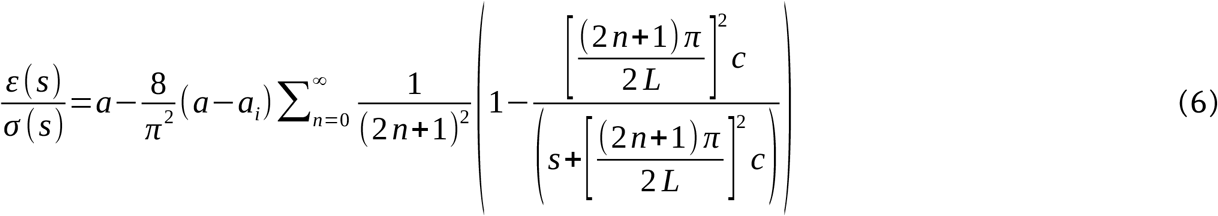

or

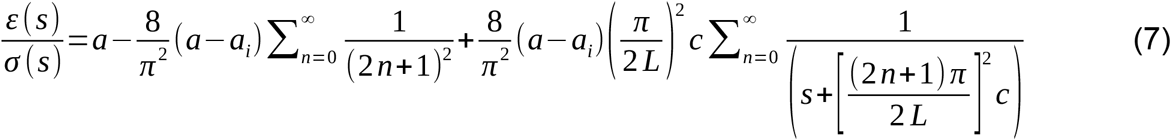

or

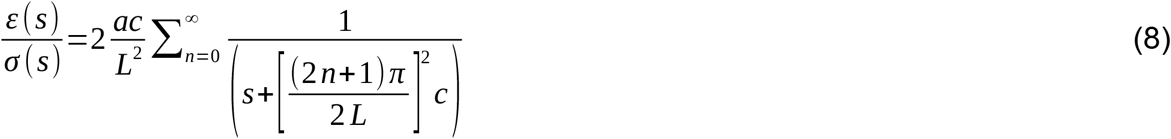

which is because

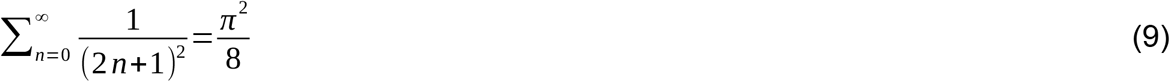

and

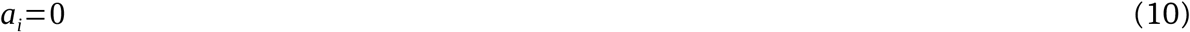

assuming fully saturated poroelastic material [20].

For a simplified approximation, *n*=0 may be assumed in Eq. (8). Accordingly:

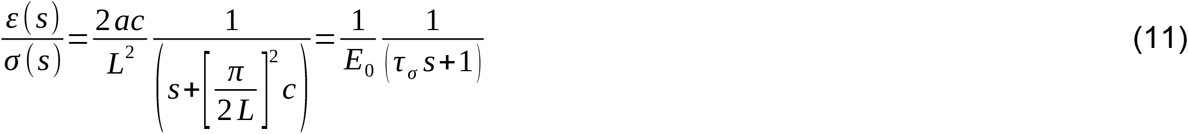

where

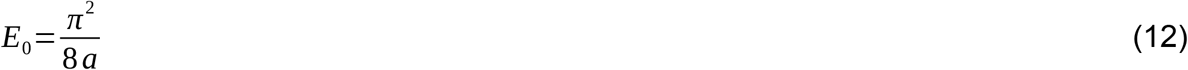

and

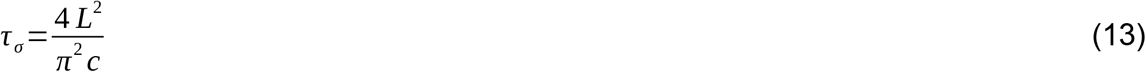

Equation (11) may be rewritten as:

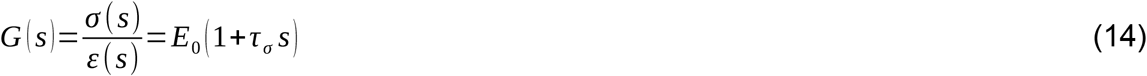

The corresponding complex modulus is

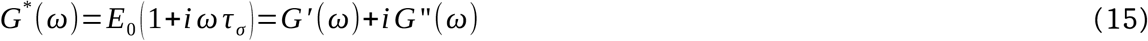

where *G*’ and *G*’’ are storage and loss modulus, which are respectively real and imaginary parts of the complex modulus *G*^*^. Therefore,

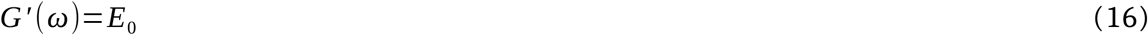

and

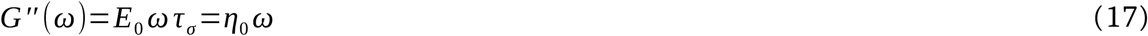

which correspond to the Kelvin–Voigt viscoelastic material of elasticity modulus *E*_0_ and viscosity *η*_0_ ([12], [21], [22]).

The continuum poroelastic column can thus be approximated as a lumped Kelvin–Voigt viscoelastic material, as represented in Fig. 1.

**Figure 1.**
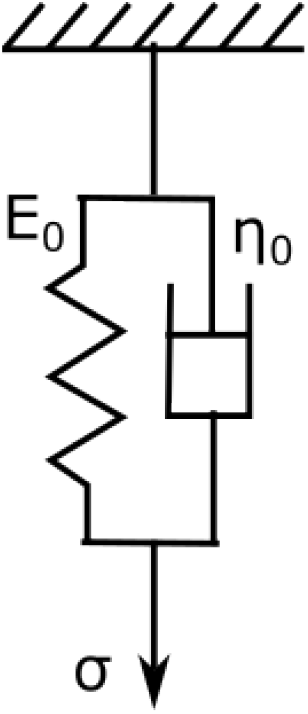
Kelvin-Voigt viscoelastic model

### 2.2. The Viscoelastic Model of Bone

The bone may be assumed to have two phases – the first is the poroelastic extracellular matrix (hereafter referred to as Phase 1, having complex modulus *E*_0_+*i η*_0_*ω*, in accordance with Section 2.1) and the other is osteocytic cell living inside these porosities (Phase 2, having complex modulus *E*_*c*_ +*iη*_*c*_*ω*). Figure 1(A) shows an idealized unit cell for the bone consisting of the solid bone matrix, fluid-filled lacunar-canalicular porosity and osteocyte living within it. The poroelastic Phase 1 (extracellular matrix) consists of the bone matrix and the fluid-filled porosities between the bone matrix and osteocytic cells (Fig. 1(B)). Phase 2, on the other hand, corresponds to the osteocytic cells. The equivalent viscoelastic model of the entire bone may be assumed to be that shown in Fig. 1(C), where *E*_*i*_ and *η*_*i*_ (with *i*=1, 2, 3) represent elastic modulus and viscosity respectively.

The parameters are related as follows (due to the fact that stiffness is proportional to the area of cross-section and modulus of elasticity, and inversely proportional to the length of the element [23]).

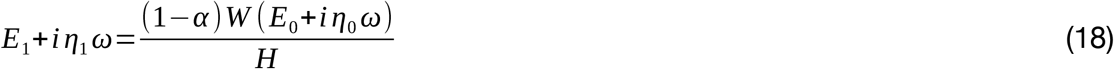

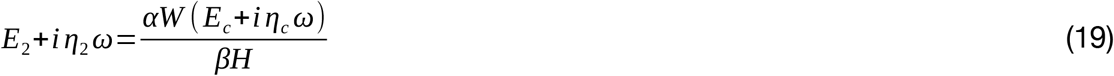

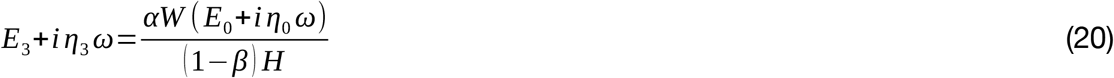

where *E*_1_ and *E*_3_ correspond to Young’s moduli of Phases 1 only, for volume fractions (1−*α*) and *α* (1−*β*) respectively; *η*_1_ and *η*_2_ are the corresponding viscosities. *E*_2_ and *η*_2_ correspond to Phase 2 with volume fraction *αβ*, where *α* (0<*α* <1) and *β* (0< *β* <1) are constants. *W* and *H* are respectively the width and the height of the unit cell.

Assuming random orientation of osteocytes in bone matrix leading to isotropic properties of bone, it may be further assumed that

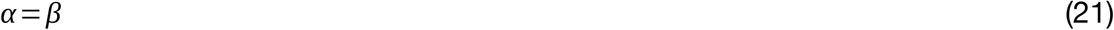

and

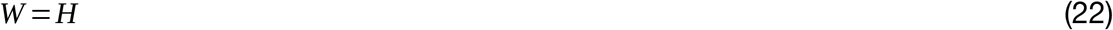

Around 10% of cortical bone volume is occupied by bone cells, about 90% of which are osteocytes [24]. Accordingly, for the 2D unit cell in consideration (Fig. 2(B)), *α* =0.3 may be assumed.

**Figure 2.**
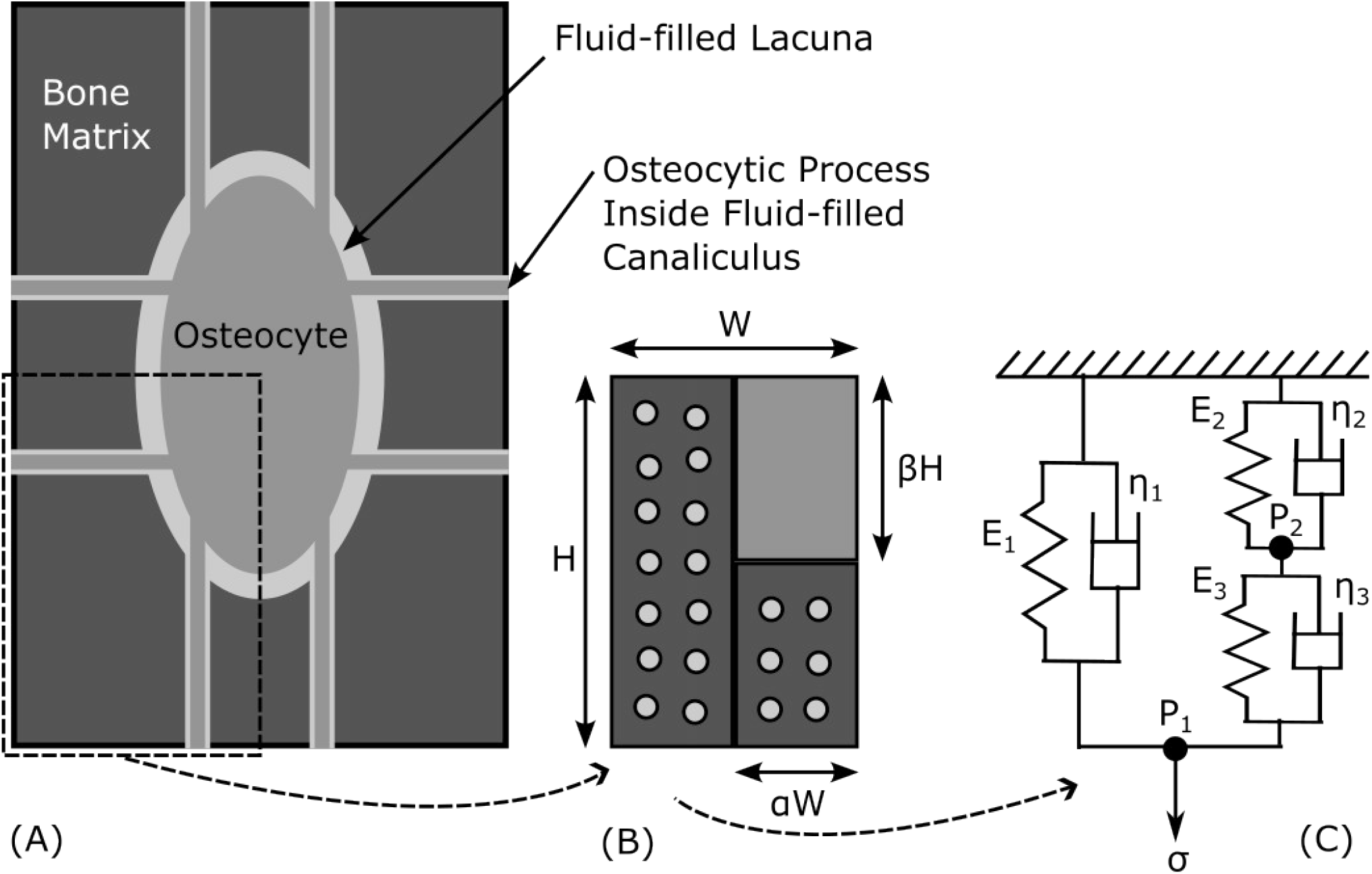
(A) Simple representation of a unit cell of the bone matrix containing fluid-filled lacuna and canaliculi inside which osteocyte along with its processes live, (B) Micromechanical two-phase representation of the bone inspired from one quarter of the unit cell (shown as dashed box in (A)), (C) Equivalent viscoelastic model.

With Eqs. (21)-(22), Eqs. (18)-(20) thus approximately become:

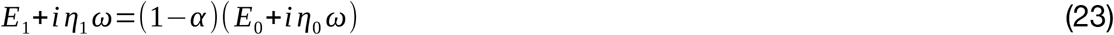

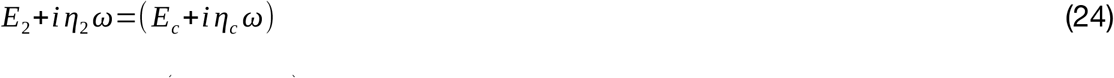

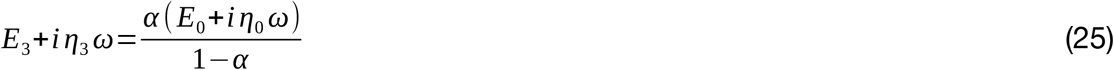

If a viscoelastic system shown in Fig. 1(C) is subjected to a stress *σ* at point P_1_ and the corresponding strain measured at that point is *ε* and that at point P_2_ is *ε*_1_, then the stress (*σ*_1_) at point P_2_ will be given by:

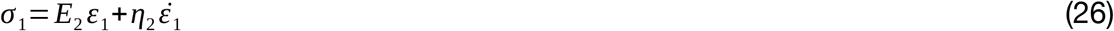

and

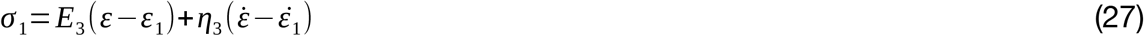

and also,

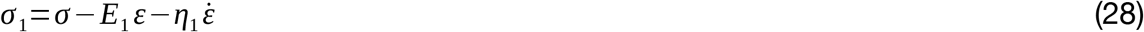

Eqs. (26), (27) and (28) respectively become the following after applying Laplace transform [25]:

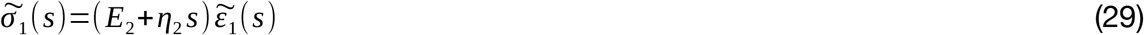

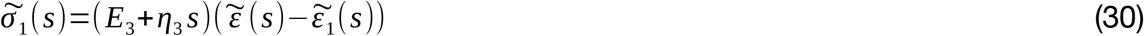

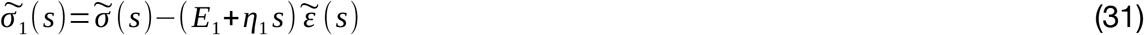

From Eqns. (29) and (30), we get:

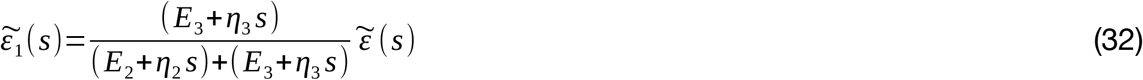

From Eqns. (29) and (32), we have

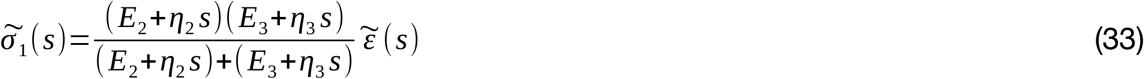

From Eqns. (31) and (33):

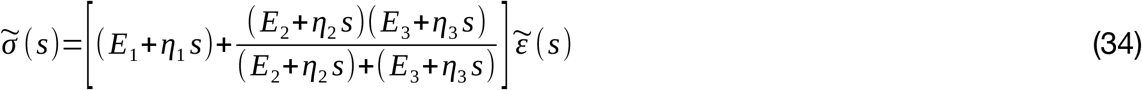

The corresponding complex modulus is given by:

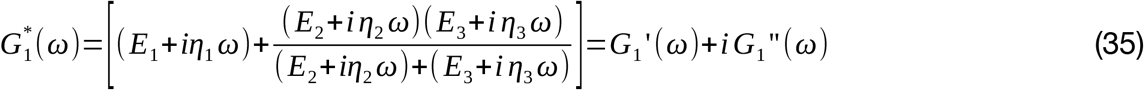

where *G*_1_’(*ω*) and *G*_1_’’(*ω*) are storage and loss modulus, respectively.

For simplicity, the following is approximately assumed, as the data in the literature roughly follow this relationship [19]:

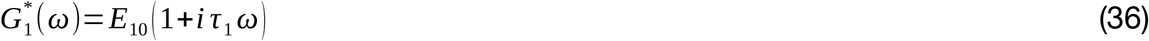

where *E*_10_ and *τ*_1_ are constants, which are chosen such as follows:

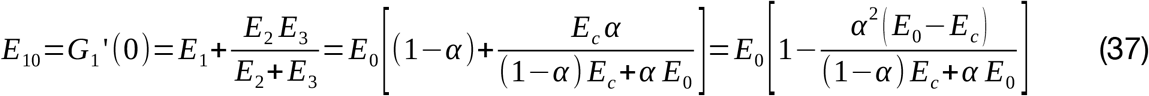

and

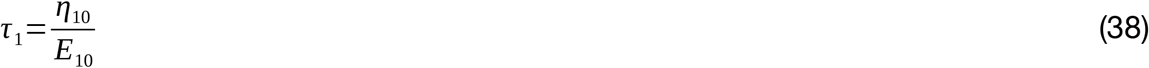

where

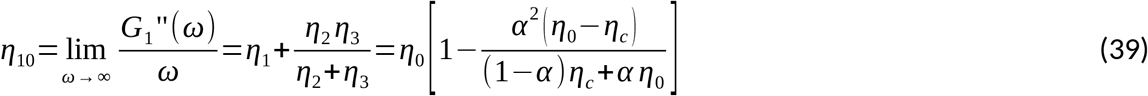

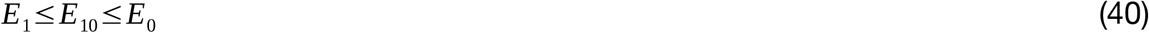

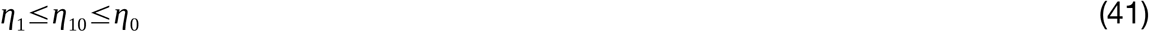

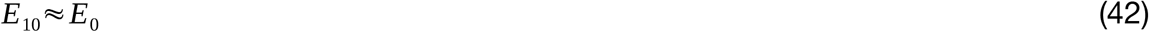

and

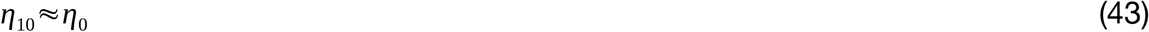

given that *α*≈0.3, *E*_*c*_ ≪ *E*_0_ and *η*_*c*_ ≪*η*_0_, viz. *E*_*c*_ ≈1×10^3^ Pa, *E*_0_ ≈1.7×10^10^ Pa, *η*_*c*_ ≈5×10^3^ Pa-s and *η*_0_ ≈1.7×10^8^, as per the literature [26], [27].

Hsieh et al (2009) presents the stress amplitude vs. strain amplitude data for female Sprague-Dawley rats of age 7–8 months old when ulna is loaded with the haversine loading waveform [19]. The corresponding experiment data may be fit to the viscoelastic model to determine the values of the constants *τ*_1_ and *E*_10_.

In response to the haversine waveform of stress:

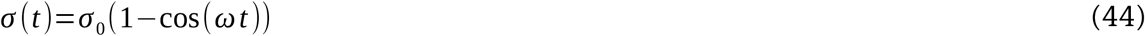

with the stress amplitude *σ*_0_> 0, the corresponding steady-state strain will be of the form:

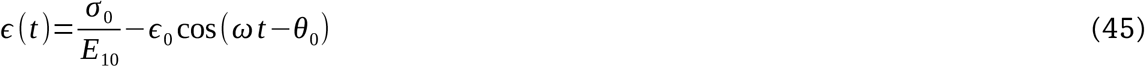

where

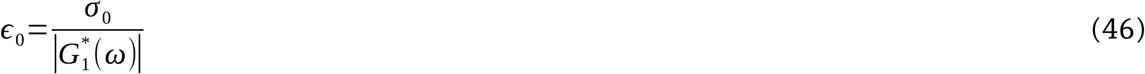

and

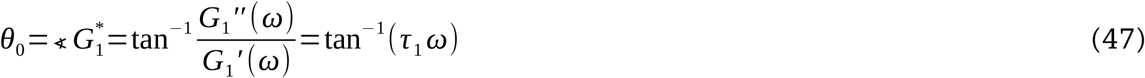

Thus, for the cyclic stress amplitude *σ*_0_, the cyclic strain amplitude is given by:

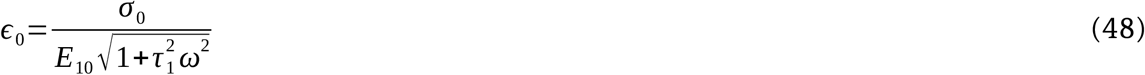

The experimentally-reported peak stress is typically the peak-to-trough amplitude of the strain. The peak strain is thus given by:

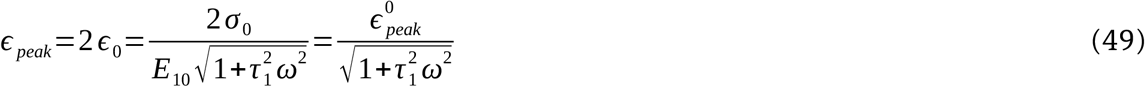

where 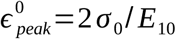 is the peak strain at zero frequency (or very-very low frequency).

The Levenberg-Marquardt Algorithm implemented through MATLAB (Mathworks Inc.) has been used for the curve fitting and determining the values of the constants *τ*_1_ and 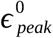 [28], [29]. The Chi-Square Goodness of Fit (i.e. Pearson’s chi-squared test) has been used to quantify the quality of curve-fitting [30].

### 2.3. Derivation of Strain Threshold

According to the First Law of Thermodynamics [31],

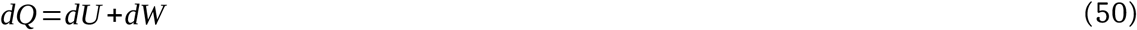

where *dQ* is the energy supplied to a system, *dU* is the change in the internal energy of the system and *dW* is the work done by the system.

For one cycle of loading,

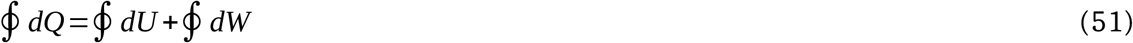

However, for a cyclic loading, the net change in the internal energy is zero, i.e.

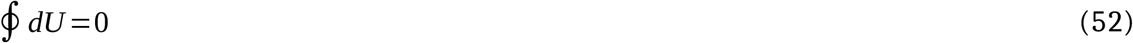

Thus,

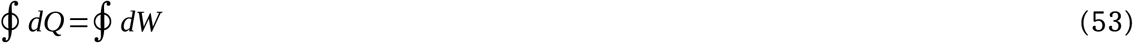

Let *ϕ* be the energy supplied to per unit volume of the loaded osteocytes (i.e. Phase 2), then

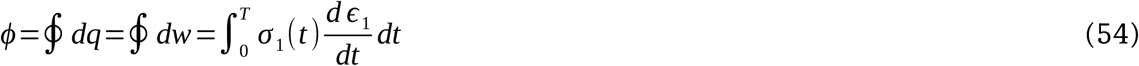

where *dq* is the energy supplied per unit volume of the system, *dw* is the work done per unit volume of the system (Phase 2) and *T* is the time period of the cyclic loading. The quantity *ϕ* is called the dissipation energy density [32].

For *σ* (*t*) and *ϵ* (*t*) given in Eqs. (44) and (45) respectively, the corresponding *ϵ*_1_(*t*) and *σ*_1_(*t*) are given by using Eqs. (32) and (33):

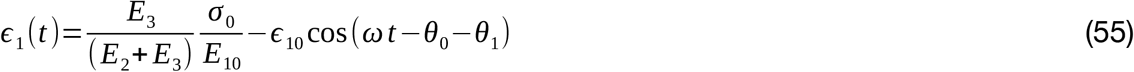

where

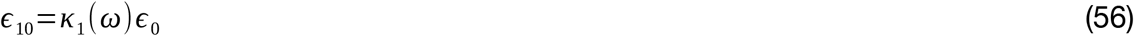

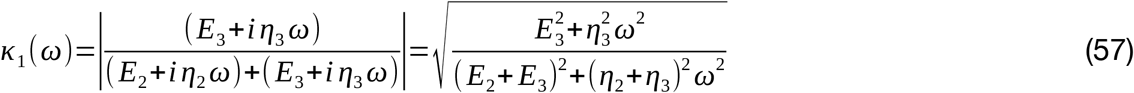

*κ*_1_ (*ω*) monotonically varies from *κ*_1_ (0)=*E*_3_ /(*E*_2_ + *E*_3_) to *κ*_1_ (∞)=*η*_3_ /(*η*_2_ +*η*_3_) as *ω* varies from 0 to ∞.

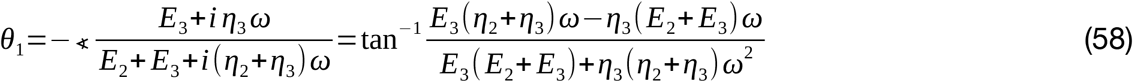

Eq. (33) can be rewritten as:

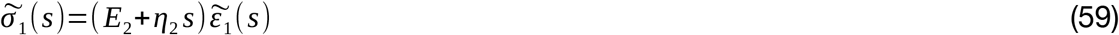

and thus

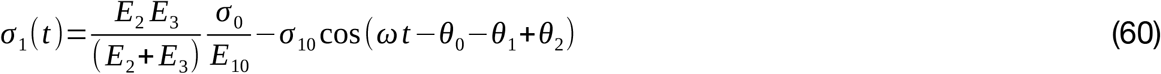

where

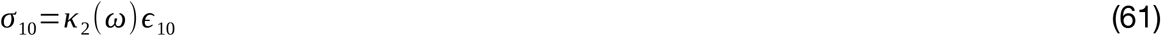

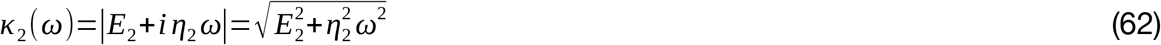

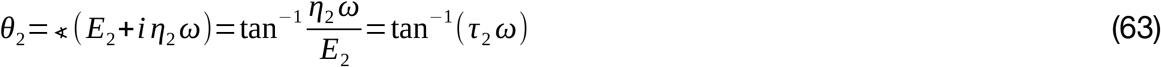

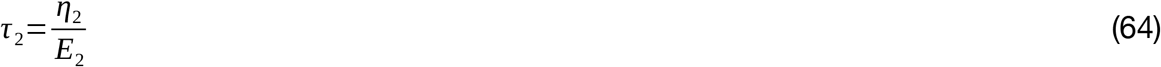

The new bone formation is assumed to depend on the work done on the osteocytes. From Eqs. (54), (55) and (60), in accordance with [32], [33], we get

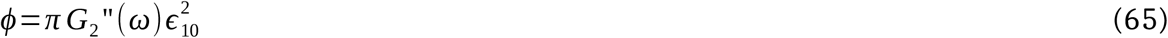

where

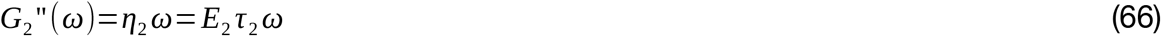

From Eqs. (56) and (65):

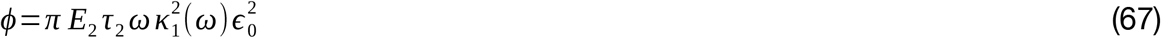

Assuming that the dissipation energy density (*ϕ*) must exceed a threshold value for new bone formation:

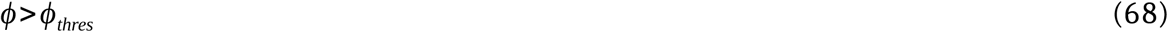

or

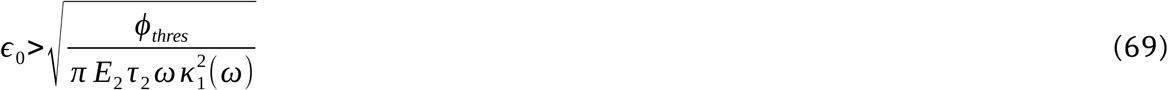

or

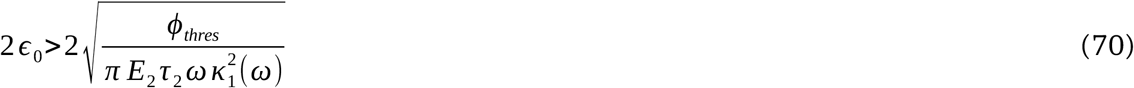

or

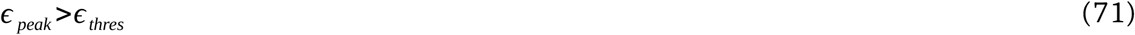

where

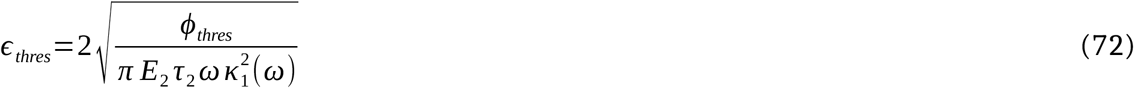

For simplicity, we will assume

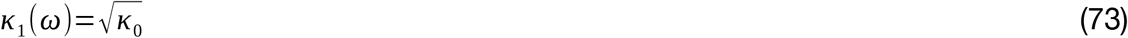

where *κ*_0_ is a constant. This assumption approximately fits the experimental data available in the literature, as shown in the results section, viz. Section 3.2.

Therefore,

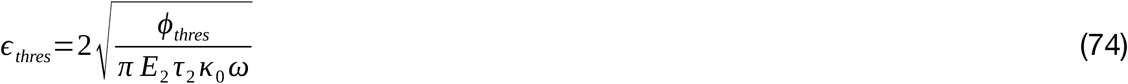

The existing experimentally determined strain threshold data (by Hsieh et al [7]) have been fitted to the derived strain threshold using the Levenberg-Marquardt Algorithm ([28], [29]) to determine the value of *ϕ*_*thres*_. MATLAB software (Mathworks Inc.) has been used for the curve-fitting.

### 2.4. Derivation of Bone Formation Rate (BFR)

It is assumed that when loading on the bones changes, the bone will do as needed (such as changing the shape and size of the bone cross-section) to attain *ϕ*_*thres*_ as the final dissipation energy density. As in Section 2.3,

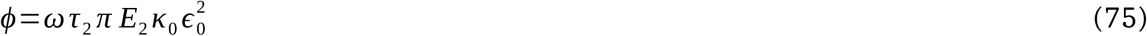

Expressing the stress *σ* in terms of axial force and induced moments [23], we get

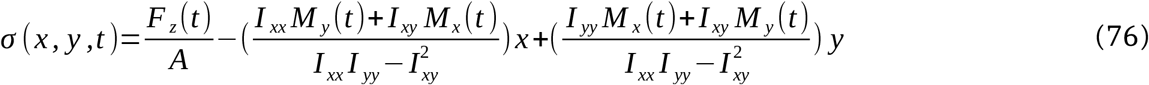

where *I*_*xx*_ and *I*_*yy*_ are second moments of the cross-sectional area of the beam about its x-axis and y-axis, respectively. *I*_*xy*_ is the corresponding product moment of area. *F*_*z*_, *M*_*x*_ and *M*_*y*_ are respectively axial force, moment about the x-axis and moment about the y-axis, respectively.

We assume the following, which is approximately true for a section in concern:

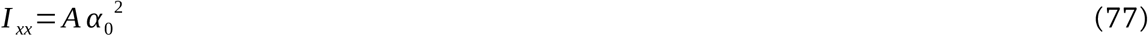

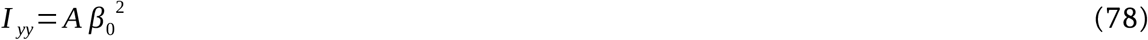

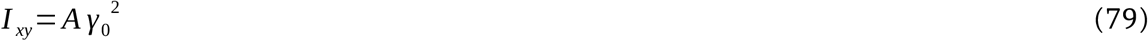

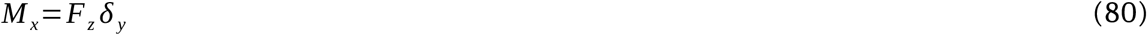

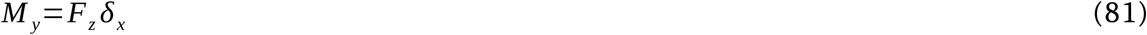

where *α*_0_, *β*_0_, *γ*_0_, *δ*_*x*_ and *δ*_*y*_ are constants.

Accordingly,

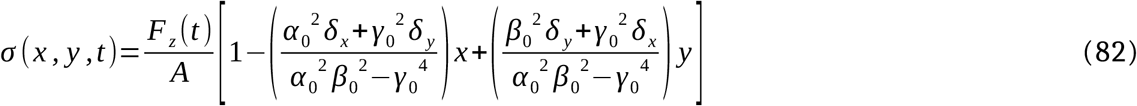

As stress *σ* is cyclic, having the haversine waveform [7], the stress amplitude is given by:

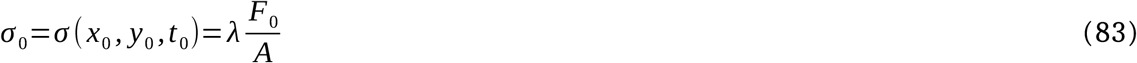

where

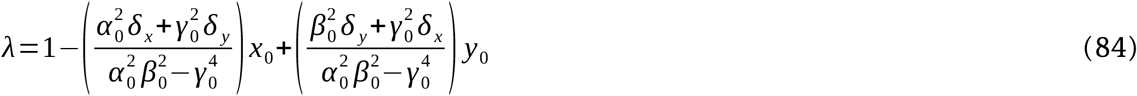

is a constant. *F*_0_ is the force amplitude of the cyclic loading. *x*_0_, *y*_0_ and *t*_0_ are the x-coordinate, y-coordinate and time corresponding to the peak stress.

From Eq. (46),

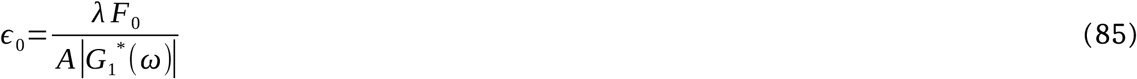

Accordingly from Eq. (75),

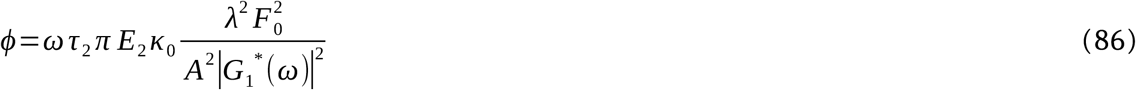

or

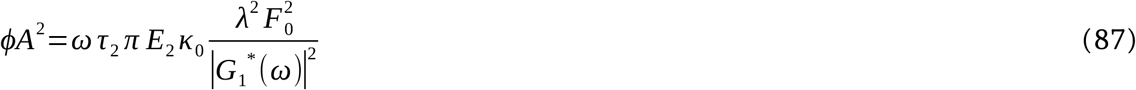

For the same cross-section and for the same loading protocol (including the number of cycles per day),

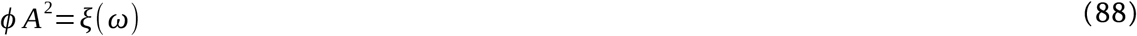

where *ξ* is a function of frequency *ω*, given by

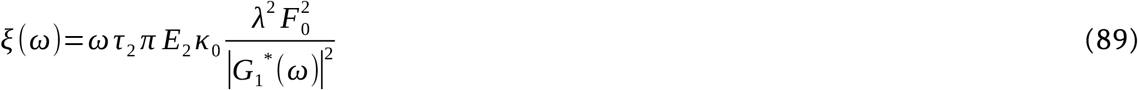

or

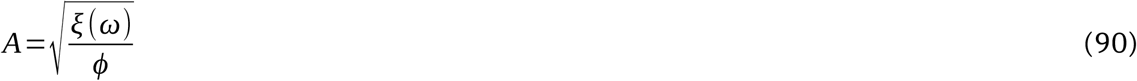

Bone Formation Rate (BFR) may be derived similar to the Mineral Apposition Rate (MAR) derived in Singh et al. [33]. Accordingly, rate of change of area is assumed to be proportional to the difference between the final area of cross-section and the current area of cross-section, i.e.

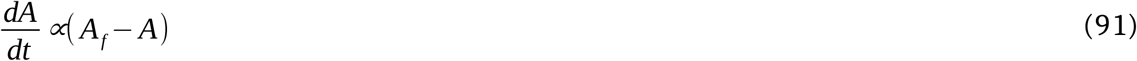

or

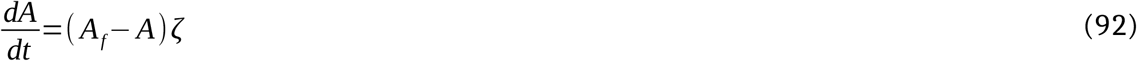

where *ζ* is the proportionality constant, which may be a function of the loading condition / protocol (including the number of cycles of loading per day). This equation may be rewritten in terms of dissipation energy density as

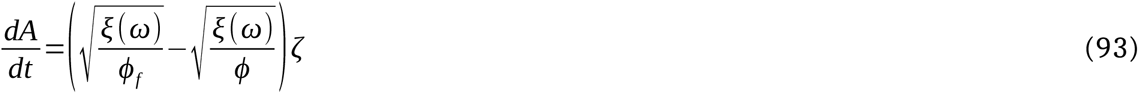

or

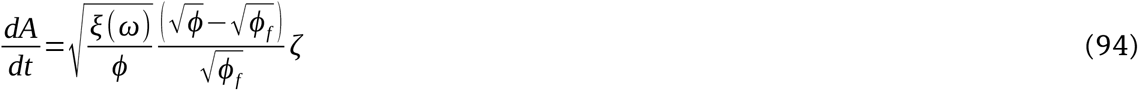

or

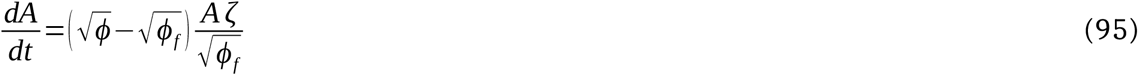

The bone formation rate (BFR) is given by,

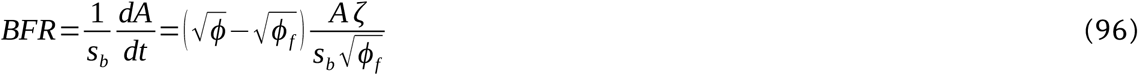

where *s*_*b*_ is the bone surface (BS), i.e. perimeter of the bone section. or,

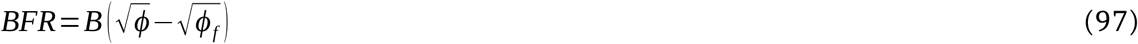

where *B* is assumed approximately to be a constant (for a slow rate of change of area, *A*) given by:

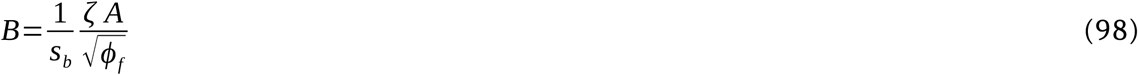

We hypothesize: *ϕ*_*f*_ =*ϕ*_*thres*_, and therefore, the BFR may be rewritten as

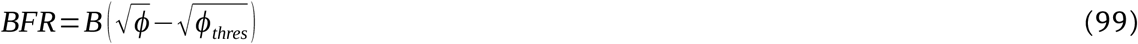

or, from Eqs. (99), (67), and (74):

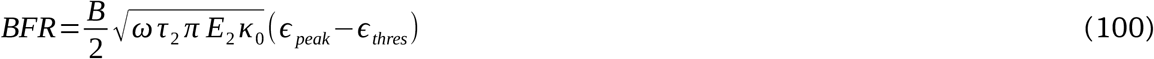

Hsieh and Turner [7] provided the experimental BFR data which varied with frequency. The Levenberg-Marquardt Algorithm ([28], [29]) has been used to fit the experimental data to the above mathematical model for BFR and accordingly determine the value of *B*.

### 2.5. Extraction of Data

Frequency versus peak strain (both tensile and compressive) data for axial loading of ulna have been approximately obtained from Fig. 2 of the work by Hsieh et al (1999) [19]. The “Preview” software (Apple Inc.) was used to get the pixel coordinates of the data, which were scaled to get the actual data using MATLAB (Mathworks Inc.). Similarly, the standard error (SE) data have also been extracted from the same source. If standard error is not specified for a particular data point, the SE (in terms of percentage) was assumed to be similar to that of the other data in the figure. Following the same methods, the Bone Formation Rate (BFR) vs. the peak strain data, along with standard errors, have been approximately obtained from Fig. 5 of the work by Hsieh and Turner [7] by using the softwares “Preview” (Apple Inc.) and MATLAB (Mathworks Inc.).

### 2.6. Statistical Analysis

Two types of statistical analysis have been carried out. Mathematical model fitted to a few given experimental values has been statistically analyzed using the “Chi Square Goodness of Fit” [30]. In addition, each experimental data point has also been compared with the corresponding value obtained from the developed mathematical model and statistically analyzed using one-sample, two-tailed Student’s t-test [34]. The MATLAB software has been used for both kinds of data analysis.

## 3. Results

### 3.1. Peak Strain as a Function of Forcing Frequency

As derived in Section 2.2,

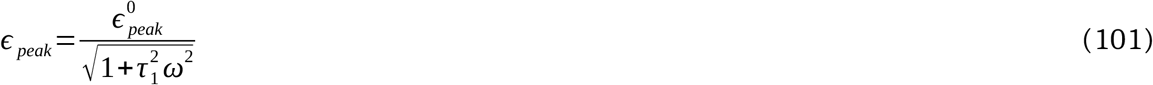

The experimental peak tensile strain values along with the respective standard errors approximately obtained from Hsieh et al (1999) [19] are tabulated in Table 1. Equation (101) is fitted with these data. Curve fitting determined the following values of parameters: 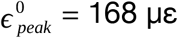 and *τ*_1_ = 0.0449 s. The peak tensile strain is plotted in Fig. 3 using a dashed line, where the experimental peak strains are shown as circles. The corresponding standard error (SE) is also shown for each of the experimental (i.e. in vivo) values. Chi Square Goodness of Fit analysis has been carried out [30]. The p-value is found to be 0.99, which indicates that the mathematical model is significantly the same as that of the experimental findings. The mathematical model’s values corresponding to the considered four forcing frequencies viz. 1, 2, 5, 10 and 20 Hz are also added in Table 1. The Student’s t-test has been carried out for each of the five frequencies to compare the experimental and model values. The corresponding p-values are also indicated in the table. As all the p-values are greater than 0.13, the model predictions are not significantly different from the experimental values.

**Table 1.** Experimental vs. model comparison of peak tensile strain in rat ulna as a function of forcing frequency; experimental values adapted from Hsieh et al (1999) [19].

| Frequency (Hz) | Experimental<br>Peak Strain ($\mu\epsilon$) | Standard Error<br>($\mu\epsilon$) | Model Peak<br>Strain ($\mu\epsilon$) | p-Value<br>(Student's t-test) |
| --- | --- | --- | --- | --- |
| 1 | 160 | 32 | 161 | 0.974 |
| 2 | 146 | 33 | 146 | 0.996 |
| 5 | 104 | 22 | 97 | 0.795 |
| 10 | 53 | 13 | 56 | 0.836 |
| 20 | 20 | 4 | 29 | 0.138 |

**Figure 3.**
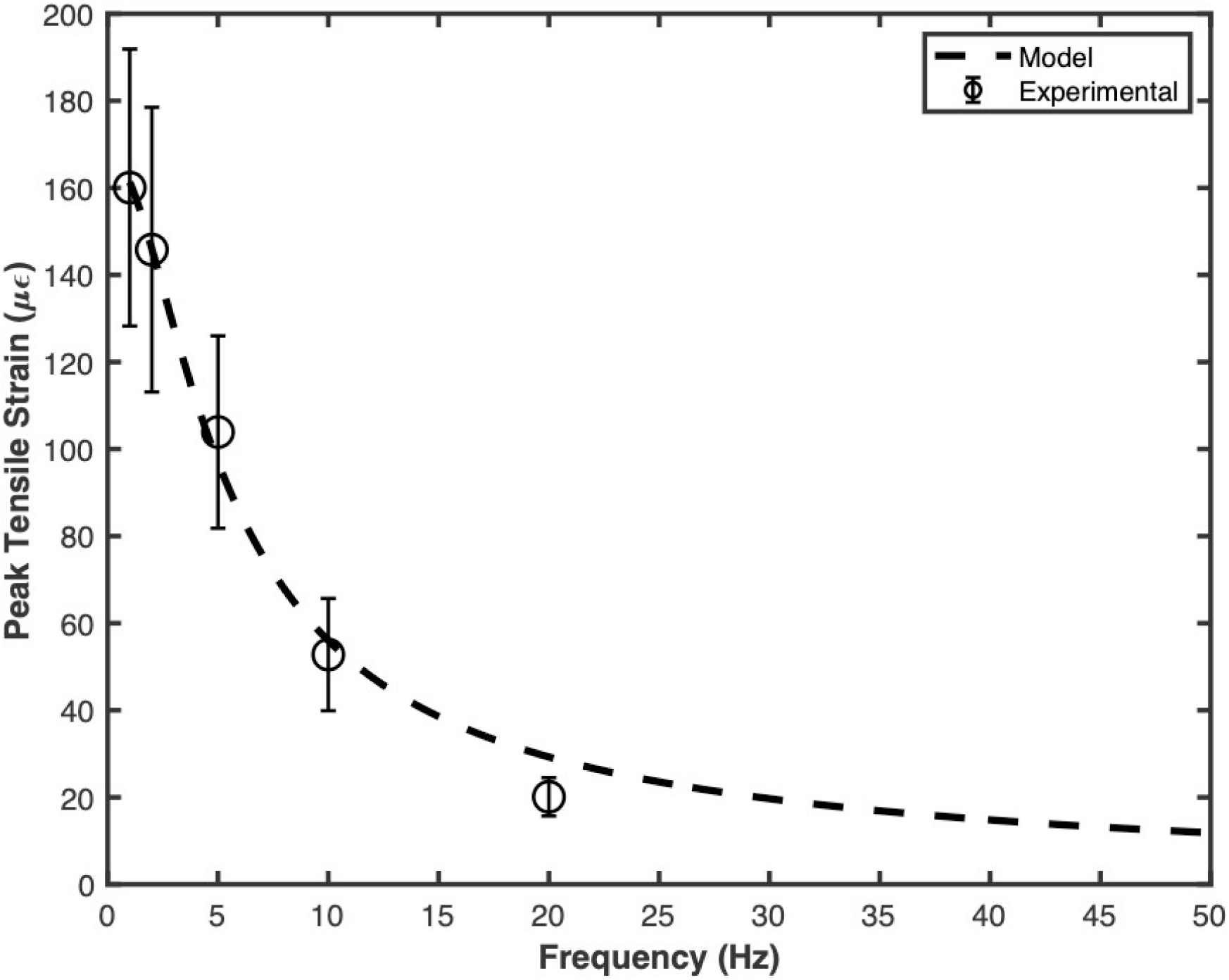
Peak tensile strain versus forcing frequency.

A similar analysis has been done for peak compressive data given by Hsieh et al (1999) [19]. The data are tabulated in Table 2 in the similar manner and plotted in Fig. 4. Curve fitting yielded the following parameters: Curve fitting determined the following values of parameters: 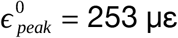 and *τ* = 0.0421 s. The p-value for the Chi Square Goodness of Fit test is 0.992. For the individual data for the five frequencies, the p-values are shown in Table 2. All p-values are more than 0.1, which indicates that the experimental and the models peak compressive strains are not significantly different.

**Table 2.** Experimental vs. model comparison of peak compressive strain in rat ulna as a function of forcing frequency; experimental values adapted from Hsieh et al (1999) [19].

| Frequency (Hz) | Experimental<br>Peak Strain ($\mu\epsilon$) | Standard Error<br>($\mu\epsilon$) | Model Peak<br>Strain ($\mu\epsilon$) | p-Value<br>(Student's t-test) |
| --- | --- | --- | --- | --- |
| 1 | -244 | 54 | -245 | 0.989 |
| 2 | -221 | 46 | -224 | 0.964 |
| 5 | -166 | 28 | -153 | 0.692 |
| 10 | -84 | 19 | -90 | 0.803 |
| 20 | -32 | 7 | -47 | 0.102 |

**Figure 4.**
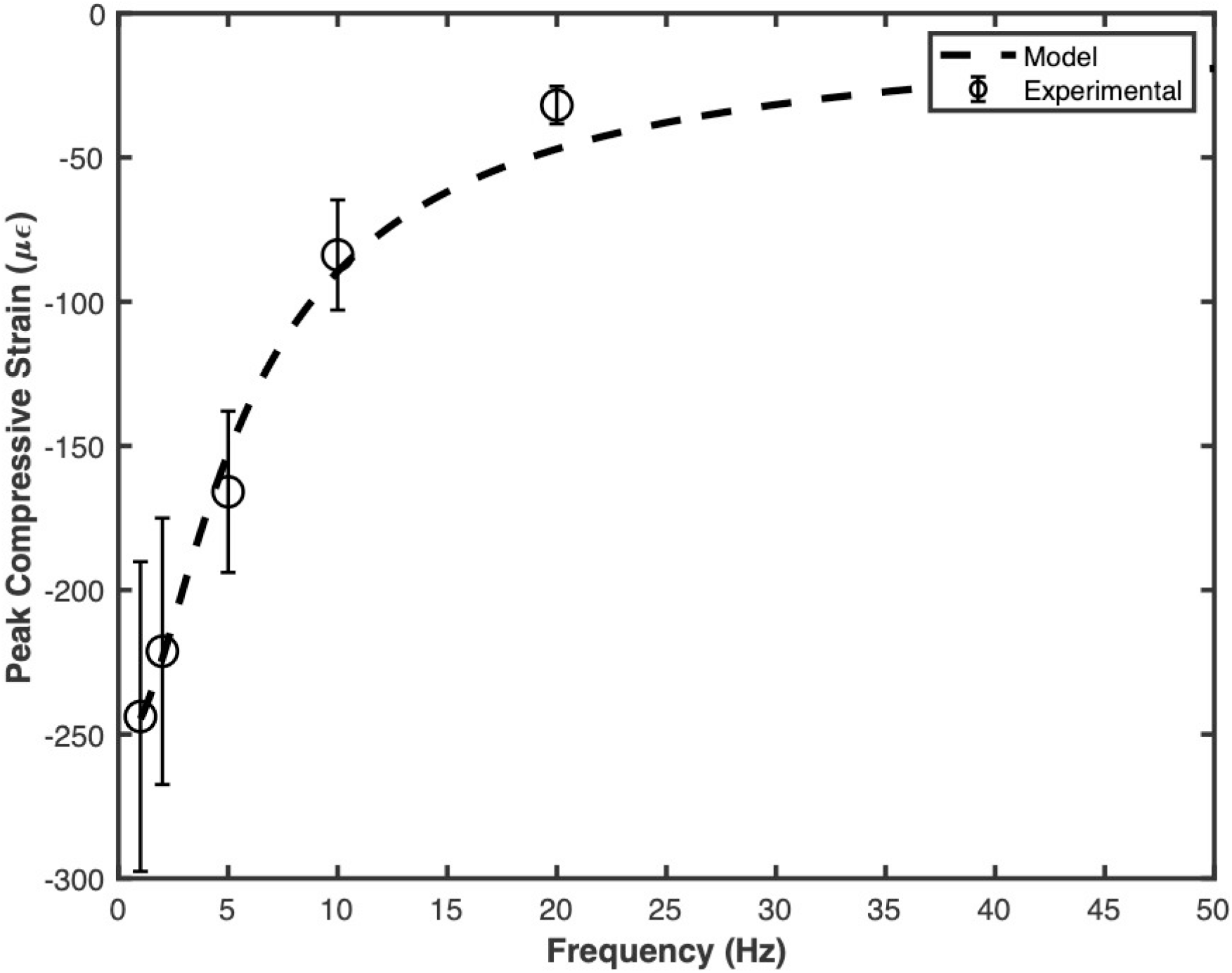
Peak compressive strain versus forcing frequency.

For both tensile and the compressive cases, *τ*_1_ values are similar, viz. 0.0449 and 0.0421 s respectively (error within 7%). It establishes that both tensile and compressive regions have the similar viscoelastic behaviour and properties.

### 3.2. Strain Threshold as a Function of Frequency

As per Section 2.3, the strain threshold is given by

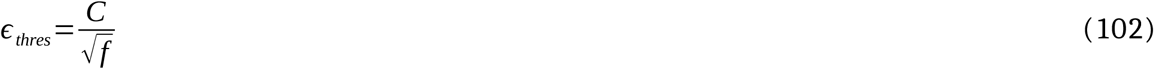

where

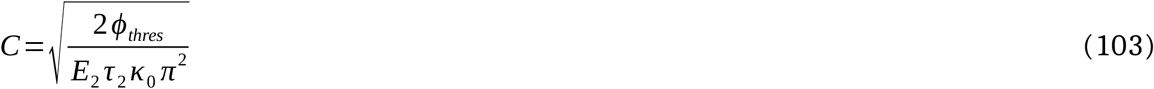

is a constant and *f* is the forcing frequency.

The experimental strain thresholds (i.e. the peak strain value at which BFR is zero) for the three frequencies (viz. 1, 5 and 10 Hz) are shown as circles in Fig. 5, which have been obtained by interpolating the BFR versus peak strain data given in Hsieh and Turner [7]. After the least squares curve-fitting, the value of *C* is found to be 1830 με/s^1/2^. The predicted strain threshold as a function of forcing frequency is also shown in the figure. The p-value for the Chi-Square Goodness of Fit test is 0.994 (i.e. p > 0.99), which means that the results of the mathematical model are not significantly different from the experimental results. Figure 5 also shows the standard error for the strain threshold estimated based on the peak-strain versus BFR plots given by Hsieh and Turner [7]. The predicted strain thresholds are thus within the standard error.

**Figure 5.**
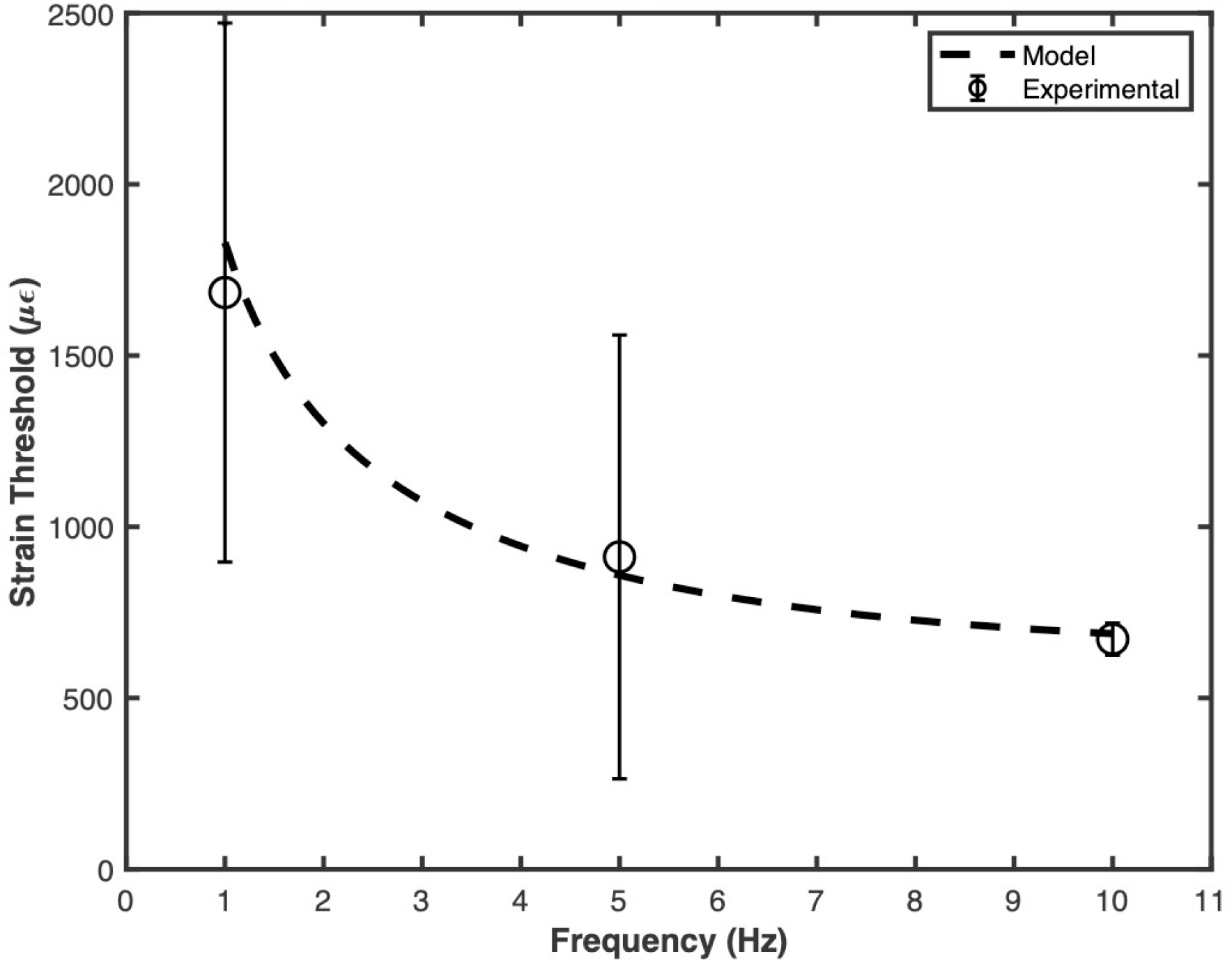
Strain threshold as a function of forcing frequency.

### 3.3. Bone Formation Rate (BFR) as a Function of Frequency

As derived in Section 2.4, the Bone Formation Rate (BFR) is given as follows:

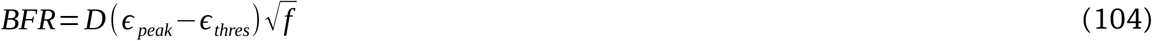

where

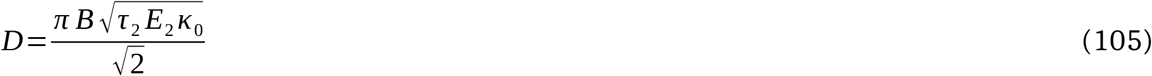

is a constant and *f* is the forcing frequency.

Equation (98) has been fitted to the in-vivo BFR vs peak-strain data in Tables 4 to 6, which are adapted approximately from Hsieh and Turner (2001) [7]. The corresponding experimental data are shown as circles, squares and triangles, respectively, for the three forcing frequencies - 1, 5 and 10 Hz. The least squares curve fitting finds the following respective values of constant *D* for the three frequencies: 0.0423, 0.0329, and 0.0773 μm-s^1/2^/με/year (i.e. mean 0.0508 ± 0.0165, standard error), which are approximately of the same order. Incorporating the corresponding value of *D*, the respective model BFRs were also computed versus the peak strain and plotted in Fig. 6.

**Table 3.** Experimental vs. model comparison of strain threshold in rat ulna as a function of forcing frequency; experimental values adapted from Hsieh and Turner (2001) [7].

| Frequency (Hz) | Experimental<br>Strain Threshold<br>($\mu\epsilon$) | Standard Error<br>($\mu\epsilon$) | Model Strain<br>Threshold ($\mu\epsilon$) | p-Value<br>(Student's t-test) |
| --- | --- | --- | --- | --- |
| 1 | 1684 | 787 | 1830 | 0.874 |
| 5 | 912 | 648 | 859 | 0.944 |
| 10 | 672 | 47 | 688 | 0.771 |

**Table 4.** Experimental vs. model comparison of bone formation rate (BFR) in rat ulna as a function of peak strain, when the forcing frequency is 1Hz (plotted in Fig. 6); experimental values adapted from Hsieh and Turner (2001) [7].

| Peak Strain ($\mu\epsilon$) | Experimental<br>Bone Formation<br>Rate ($\mu\text{m}/\text{year}$) | Standard Error<br>($\mu\epsilon$) | Model Bone<br>Formation Rate<br>($\mu\text{m}/\text{year}$) | p-Value<br>(Student's t-test) |
| --- | --- | --- | --- | --- |
| 1643 | -0.4 | 19 | -7.9 | 0.732 |
| 2767 | 33.1 | 32 | 39.6 | 0.864 |
| 3885 | 93.5 | 39 | 86.8 | 0.885 |
| 5009 | 132.4 | 68 | 134.3 | 0.980 |

**Table 5.** Experimental vs. model comparison of bone formation rate (BFR) in rat ulna as a function of peak strain, when the forcing frequency is 5Hz (plotted in Fig. 6); experimental values adapted from Hsieh and Turner (2001) [7].

| Peak Strain ($\mu\epsilon$) | Experimental<br>Bone Formation<br>Rate ($\mu\text{m}/\text{year}$) | Standard Error<br>($\mu\epsilon$) | Model Bone<br>Formation Rate<br>($\mu\text{m}/\text{year}$) | p-Value<br>(Student's t-test) |
| --- | --- | --- | --- | --- |
| 1063 | 5.0 | 17 | 15.1 | 0.600 |
| 1799 | 24.4 | 19 | 69.2 | 0.083 |
| 2515 | 147.8 | 72 | 121.8 | 0.757 |
| 3244 | 176.0 | 90 | 175.5 | 0.996 |

**Table 6.** Experimental vs. model comparison of bone formation rate (BFR) in rat ulna as a function of peak strain, when the forcing frequency is 10 Hz (plotted in Fig. 6); experimental values adapted from Hsieh and Turner (2001) [7].

| Peak Strain ($\mu\epsilon$) | Experimental<br>Bone Formation<br>Rate ($\mu\text{m}/\text{year}$) | Standard Error<br>($\mu\epsilon$) | Model Bone<br>Formation Rate<br>($\mu\text{m}/\text{year}$) | p-Value<br>(Student's t-test) |
| --- | --- | --- | --- | --- |
| 552 | -25.2 | 8 | -33.2 | 0.376 |
| 900 | 45.9 | 11 | 51.7 | 0.642 |
| 1281 | 214.2 | 51 | 145.0 | 0.251 |
| 1629 | 188.7 | 39 | 229.9 | 0.354 |

**Figure 6.**
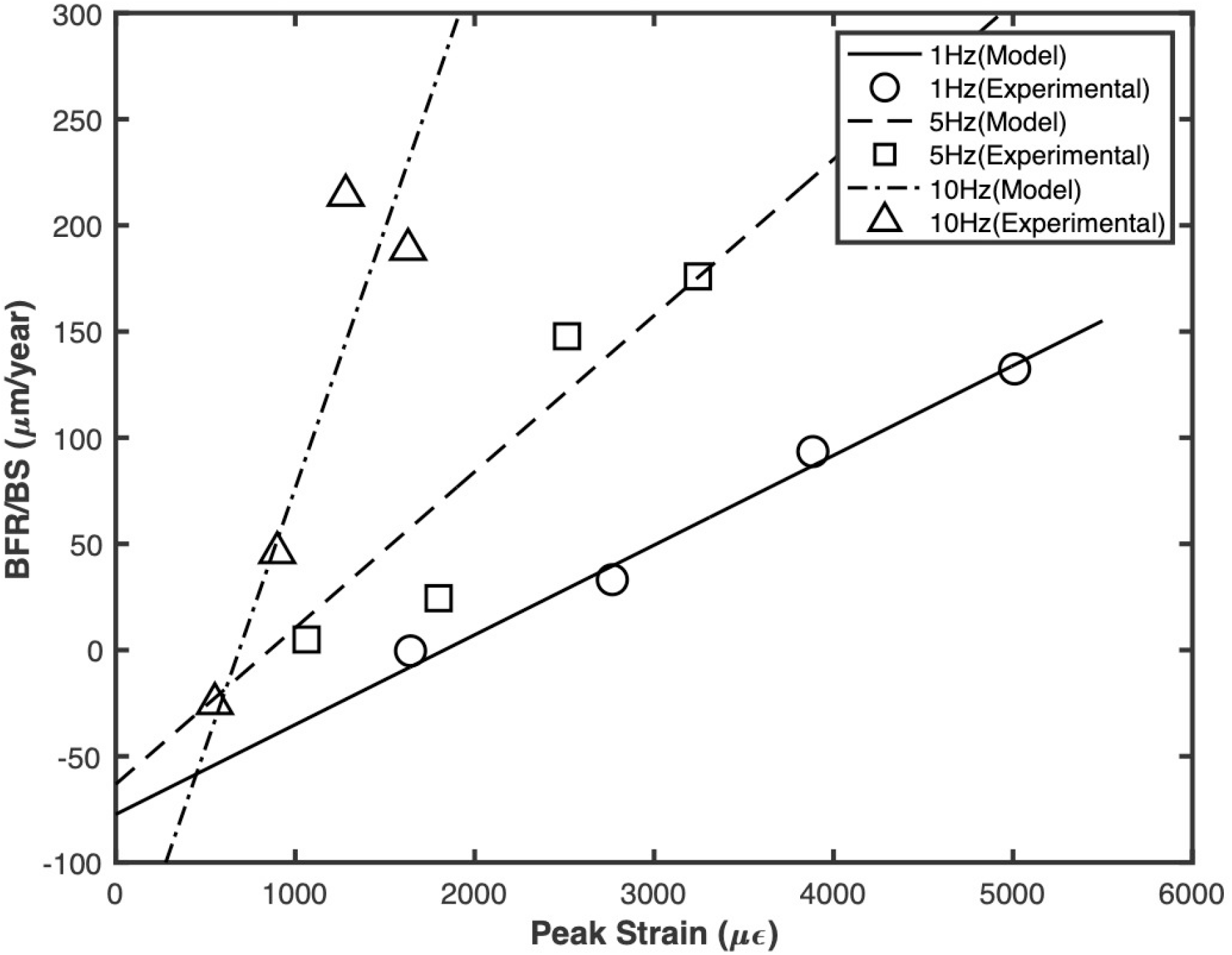
Bone Formation Rate (BFR/BS) as a function of peak strain.

The BFR predicted by the mathematical model is shown in Fig. 6 as a solid, dashed and black dash-dotted lines, respectively, for the forcing frequencies 1 Hz, 5 Hz and 10 Hz. The p-values for the Chi-Square Goodness of Fit test are 0.98, 0.06 and 0.92, respectively, which (all p>0.05) mean that the mathematical model prediction is not significantly different from the experimental values.

The standard error data adapted from Hsieh and Turner [7] are respectively given in Tables 4, 5, 6 and shown respectively in Figs. 7, 8 and 9 for the three respective forcing frequencies 1, 5 and 10 Hz.

**Figure 7.**
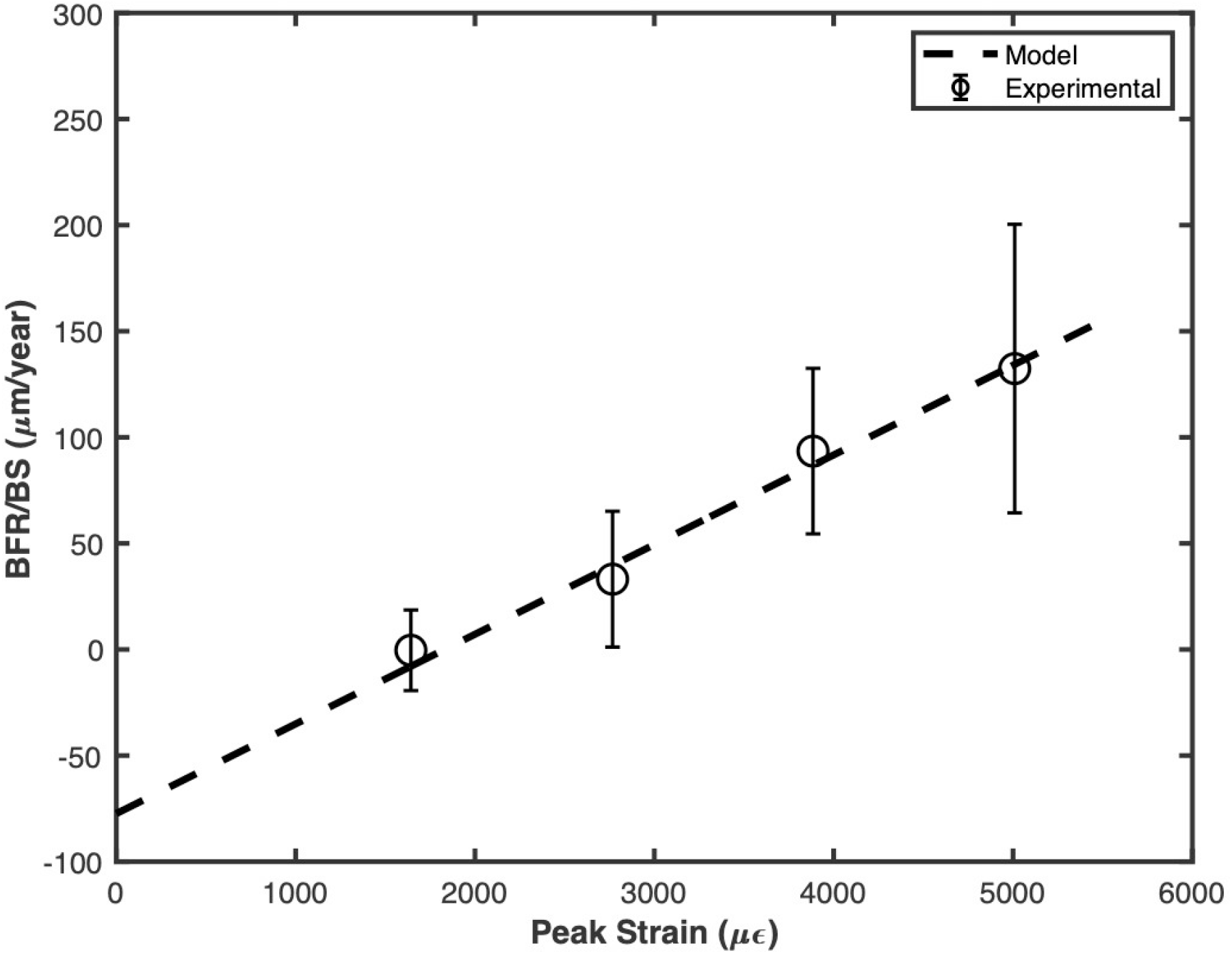
Bone Formation Rate (BFR/BS) as a function of peak strain for the frequency 1 Hz. Standard error (SE) for each of the experimental data is also shown.

**Figure 8.**
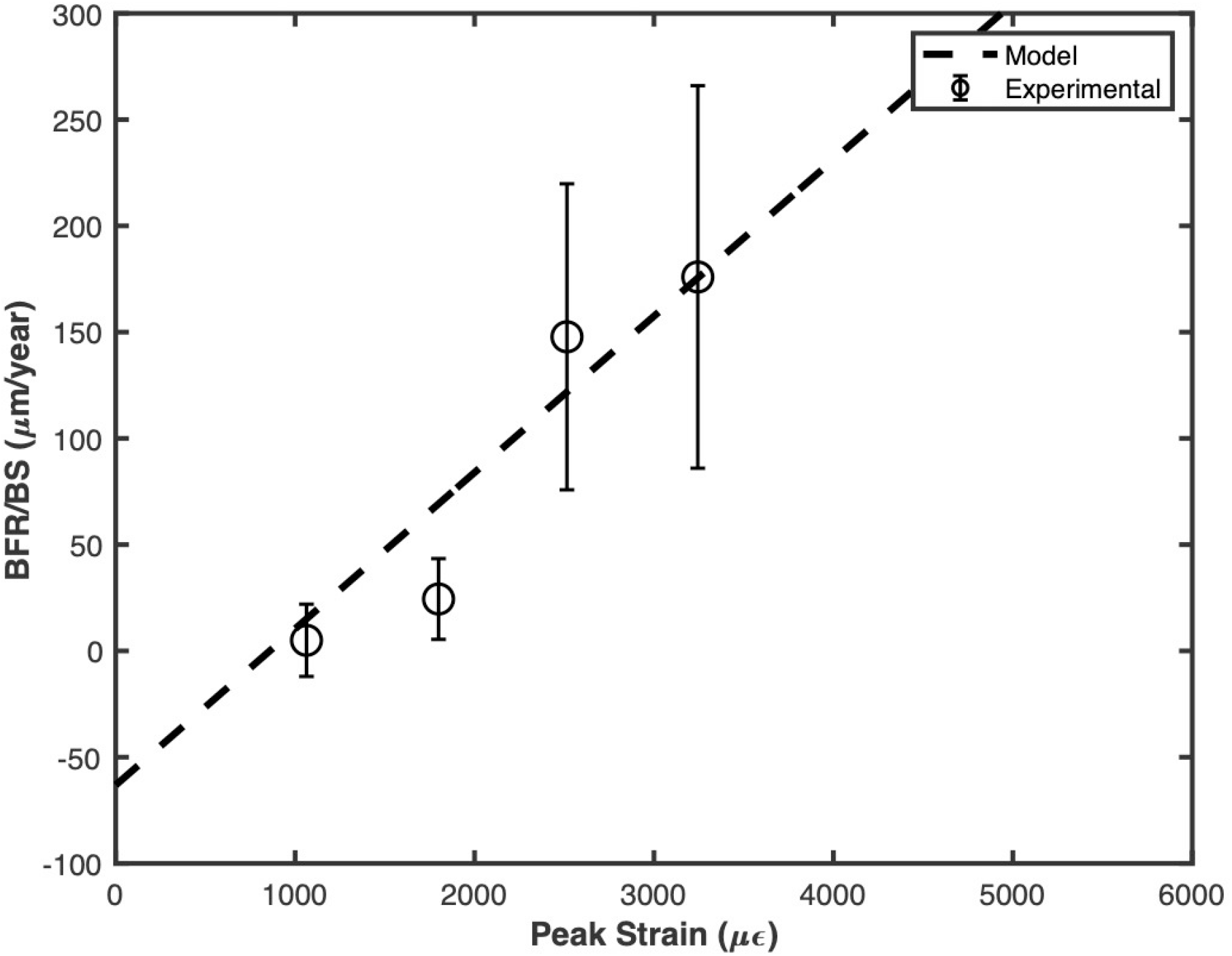
Bone Formation Rate (BFR/BS) as a function of peak strain for the 5Hz forcing frequency. Error bars indicate standard error (SE).

**Figure 9.**
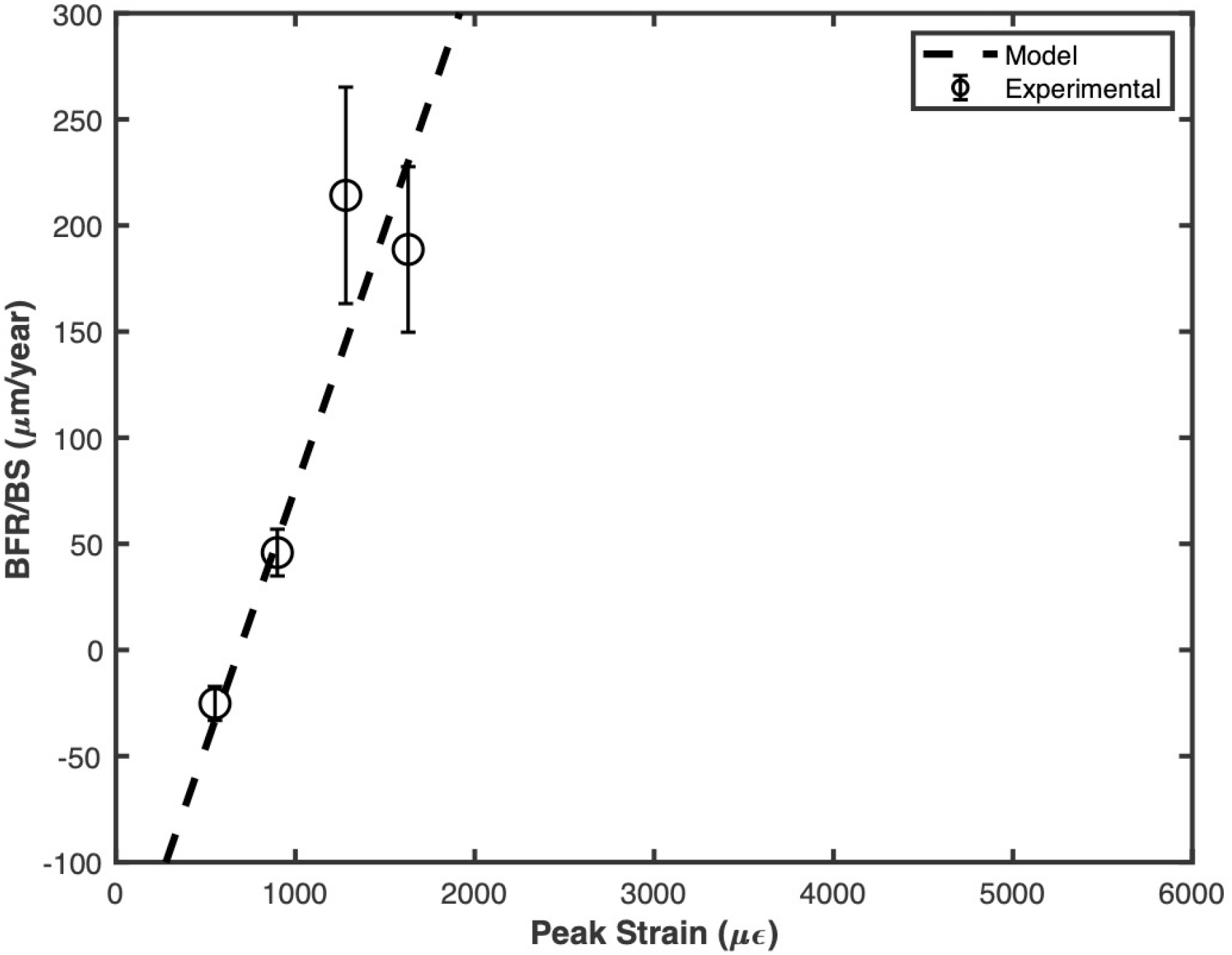
Bone Formation Rate (BFR/BS) as a function of peak strain for the 10 Hz frequency (corresponding standard errors indicated by error bars).

Student’s t-test (one-sample, two-tailed) was conducted to compare each of the experimental BFR to the corresponding value predicted by the developed model. The corresponding p-values are given in Tables 4, 5 and 6 respectively for the three frequencies, viz. 1Hz, 5Hz and 10Hz. In all cases, p > 0.05 which means that the model predictions are not significantly different from the corresponding experimental values.

### 3.4. Prediction of Strain Threshold and BFR

Hsieh et al. [8] used the similar loading protocol and animals as that used by Hsieh and Turner [7], except that the frequency for the haversine waveform was 2Hz in the case of Hsieh et al. [8]. For 2 Hz loading, the predicted strain threshold as per Eq. (102) is 1294 με, which is not significantly different from the experimental strain threshold of 1331 με (p=0.96, Student’s t-test). This threshold was calculated by the linear regression analysis of data given in Table 7, which was in turn adapted from the work of Hsieh et al. [8] for the midsection, leaving the data related to a higher strain, which resulted into fracture of the loaded bone. Note that in this statistical analysis, a standard error (SE) of 648 was assumed for 2Hz data (i.e. the same as that for 5Hz data in Table 3 and less than that for 1Hz data in the same table).

**Table 7.**
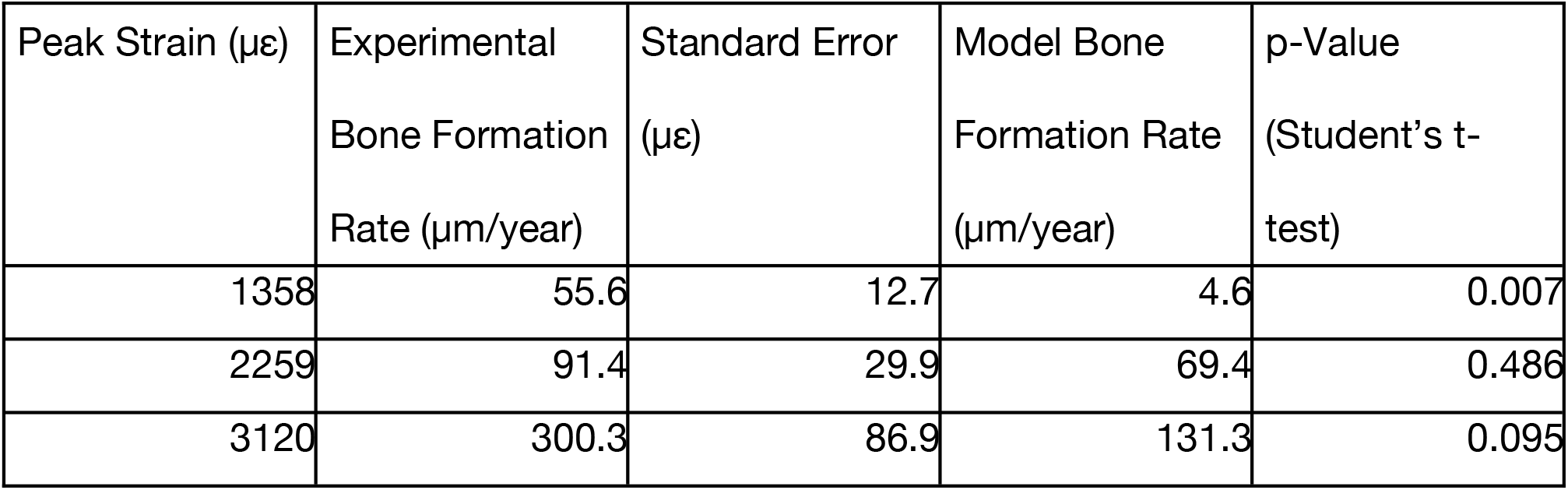
Experimental vs. model comparison of bone formation rate (BFR) in rat ulna as a function of peak strain, when the forcing frequency is 2Hz (plotted in Fig. 10); experimental values adapted from Hsieh et al. (2001) [8].

| Peak Strain ($\mu\epsilon$) | Experimental<br>Bone Formation<br>Rate ($\mu\text{m}/\text{year}$) | Standard Error<br>($\mu\epsilon$) | Model Bone<br>Formation Rate<br>($\mu\text{m}/\text{year}$) | p-Value<br>(Student's t-test) |
| --- | --- | --- | --- | --- |
| 1358 | 55.6 | 12.7 | 4.6 | 0.007 |
| 2259 | 91.4 | 29.9 | 69.4 | 0.486 |
| 3120 | 300.3 | 86.9 | 131.3 | 0.095 |

The BFR has been predicted for the three peak strains for the loading frequency of 2Hz using the mean value of *D* obtained in Section 3.3, viz. *D* = 0.0508 μm-s^1/2^/με/year. The corresponding BFR values are plotted in Fig. 10 and shown in Table 7, along with the corresponding p-values for Student’s t-test. Except for the first peak strain, i.e. 1358 με (which is close to the threshold strain), the p-values are more than 0.05. This suggests that the model BFR values are not significantly different from the experimental values. The p-value for the ‘Chi-Square Goodness of Fit’ test is 0.55, which indicates that the model results are not significantly different from the experimental values. The value of parameter *D* obtained by the linear regression analysis of experimental data in Table 7 is 0.098 μm-s^1/2^/με (which is the slope of the solid line in Fig. 8), which is not significantly different from the model mean value of *D* = 0.0508 ± 0.0165 μm-s^1/2^/με/year (p = 0.109, t-test).

**Figure 10.**
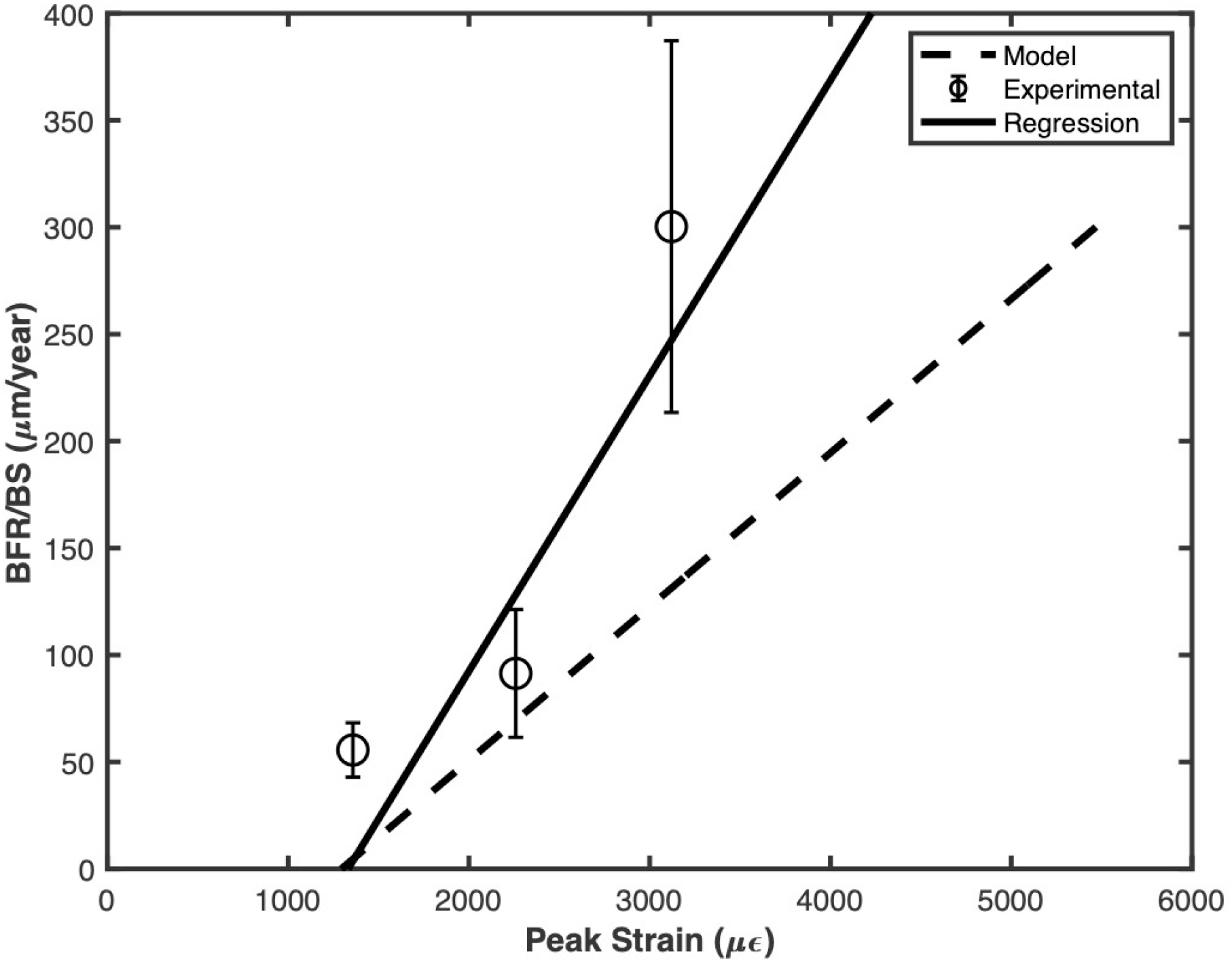
Bone Formation Rate (BFR/BS) as a function of peak strain for the 2Hz frequency (corresponding standard errors indicated by error bars).

## 4. Discussions

### 4.1. BFR as a Function of Frequency at a Constant Force Magnitude

Warden and Turner presented BFR values for mouse ulna as a function of frequency, when the load magnitude is kept constant [35]. They particularly observed that the BFR first increases with frequency and then decreases with frequency. The present work approximately agrees with that observation.

As per Eq. (99),

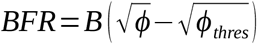

where (as per Eqs. (86), (36) and (103), respectively)

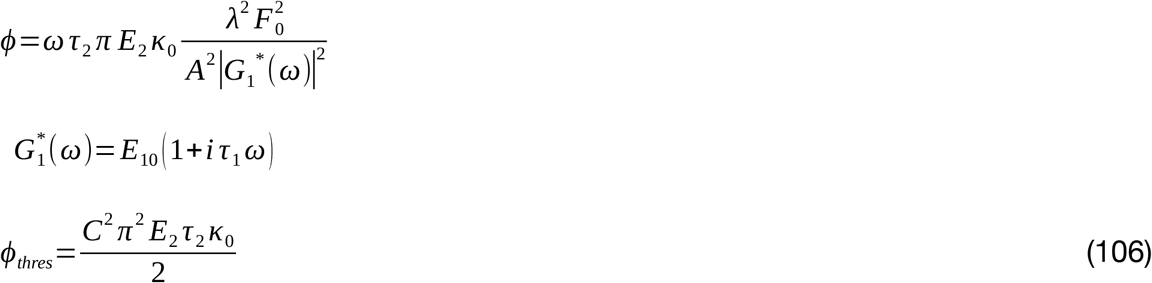

The dissipation energy density is thus given by

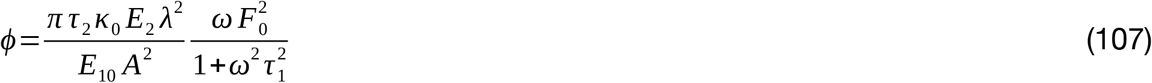

or

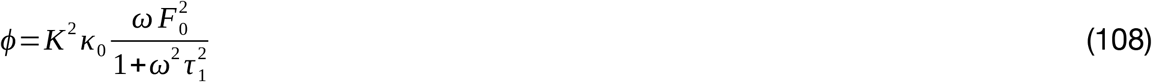

where

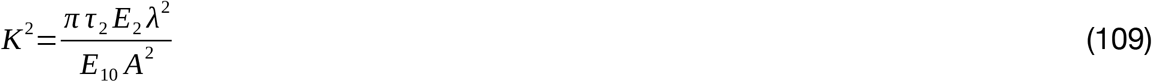

BFR is thus given by

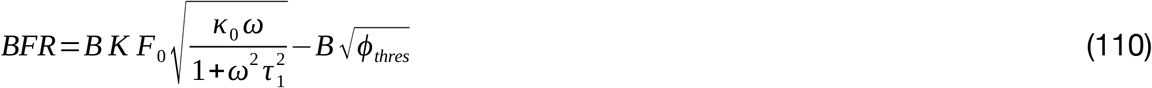

From (57) and (73),

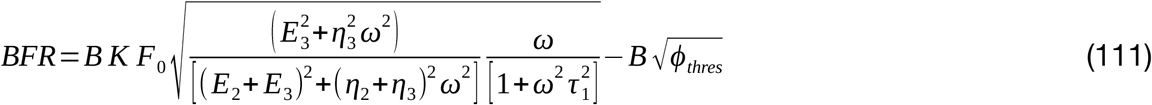

or

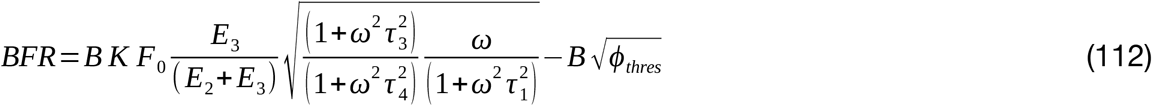

or

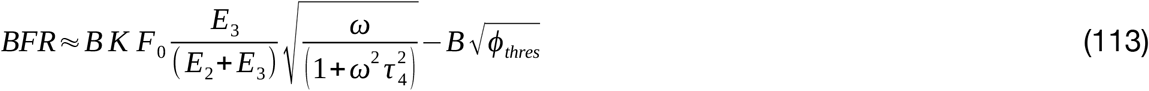

where

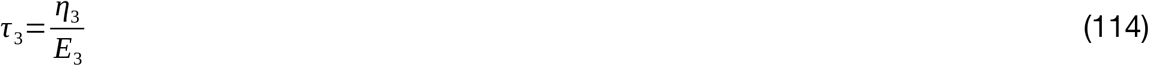

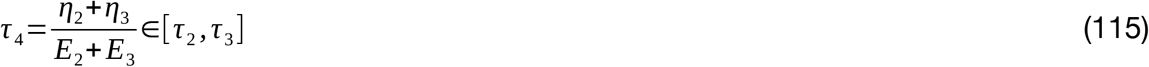

as from Eqs. (23)-(25) and (37)-(43):

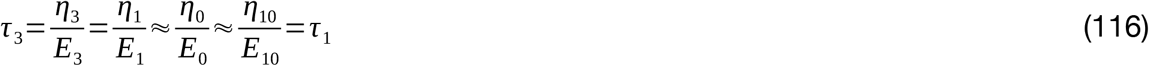

As per Eq. (104), BFR is a function of frequency and force amplitude *F*_0_. Theoretically, for a constant force amplitude, the BFR is maximum where *ω*=1/ *τ*_1_, or *f* =1/(2 *π τ*_1_). As per Section 3.1, *τ*_1_ = 0.0421 s, approximately, which predicts that at a constant load amplitude, BFR is maximum at frequency *f* = 3.78 Hz. The BFR increases with the frequency until this frequency and the BFR decreases with frequency after this frequency. However, in the case of Warden and Turner [35], the frequency at which the peak BFR happens is after 5 Hz. The title of the work itself says that “Mechanotransduction in cortical bone is most efficient at loading frequencies of 5-10 Hz”.

Equation (77) has been fit to the data reported by Warden and Turner [35]. The computed optimal model parameters are as follows: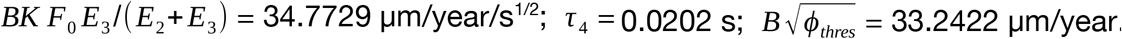. Figure 11 compares the model values to the experimental values. The fitted curve is not significantly different from the experimental data (*p* = 0.996, chi square goodness of fit test). Equation (113) thus fits the data considerably well.

**Figure 11.**
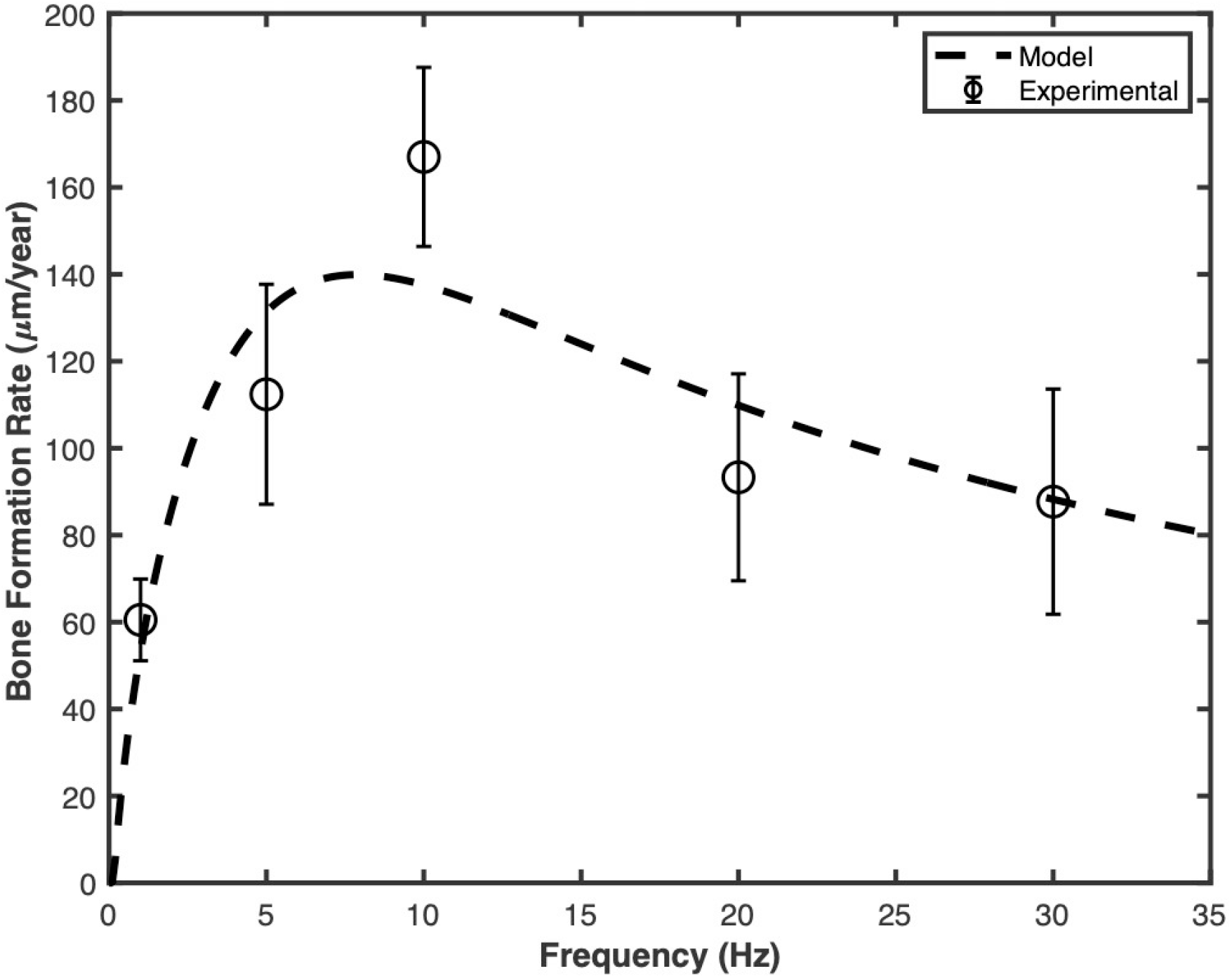
Bone Formation Rate (BFR/BS) as a function of loading frequency at a constant load magnitude (corresponding standard errors indicated by error bars).

### 4.2. Biological Basis of Dissipation Energy Density

Calcium signaling is believed to be a key mediator in bone mechanotransduction [36]; there are already some existing bone adaptation models based on calcium signalling ([11], [12]). It has been experimentally observed that there is a considerable oscillations in cytoplasmic calcium concentration as a response to tissue-level mechanical loading [36]. Jing et al. found the percentage of osteocytes responding to mechanical loading (via intracellular calcium oscillation) to be directly proportional to the loading magnitude [37]. Lewis et al., however, found the proportionality constant to be non-monotonically varying with frequency, viz. first increasing with frequency and then decreasing with frequency, when the loading on the bone tissue is kept constant [38]. The number of load-responsive osteocytes (as a percentage of the total number of osteocytes) per unit stress is shown in Fig. 11, where circles represent the data points and the error bars represent the standard error. This trend is akin to the dissipation energy density’s variation with frequency at a constant stress, as expressed in Eq. (113).

From Eqs. (23), (25) and (108), we get

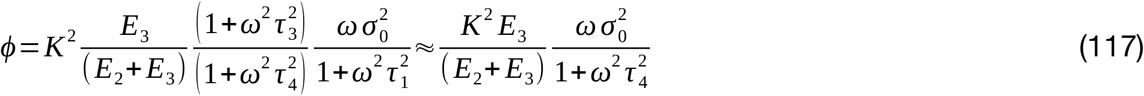

We accordingly hypothesise that the percentage (*p*_*Ca*_) of osteocytes responsive to loading is given by

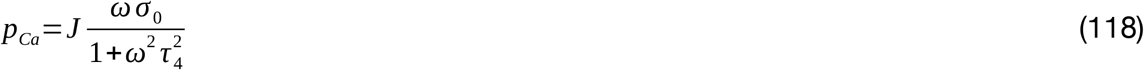

where *J* is a constant.

Equation (118) has been fitted to the experimental data in Fig. 12 and the following optimal parameters have been obtained: *J* = 0.5416 mm^2^ s / N and *τ*_4_ = 0.2145 s. The fitted model is shown as the dashed line in Fig. 12. The fitted model values are not significantly different from the experimental values (p = 0.817, chi square goodness of fit test).

**Figure 12.**
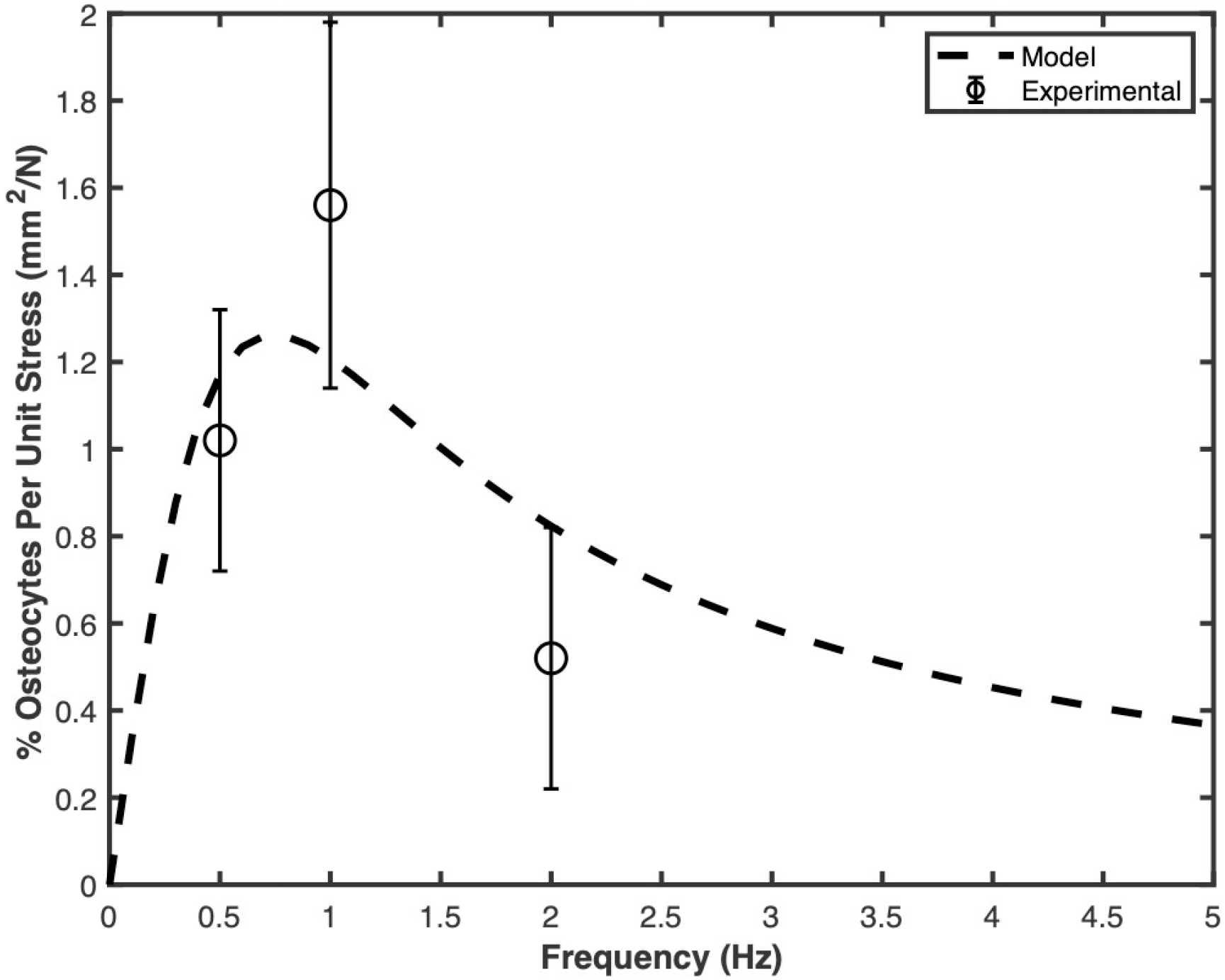
Percentage of osteocytes responding to per unit tissue-level stress magnitude as a function loading frequency (corresponding standard errors indicated by error bars). Data adapted from Lewis et al. [38].

Jing et al. also reported the number of calcium spikes to be approximately linearly varying with load magnitude [37]. Based on the data presented by Jing et al. [37], increase in calcium spikes (with respect to no exogenous loading) per unit stress is 0.171 ± 0.008 mm^2^/N, where the error represents the standard error. Accordingly, the increase in calcium spikes (*n*_*Ca*_) may be assumed to be proportional to the stress applied, that is

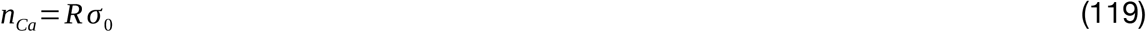

where *R* is a constant.

Equation (80) fits very well to the data presented by Jing et al. [37] with *R* = 0.171 mm^2^/N (p = 0.993 for the chi square goodness of fit).

Combining equations (118) and (119),

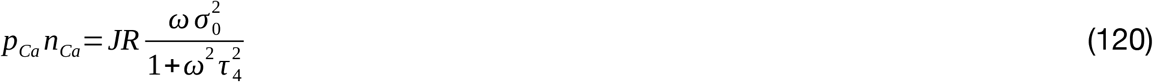

From equations (117) and (120),

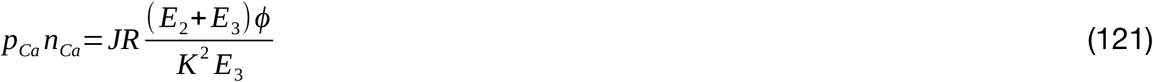

which simply states that

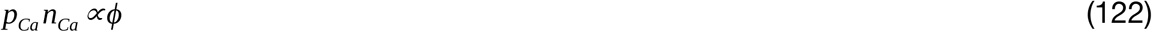

In other words, the quantity of calcium oscillation (viz. percentage (*p*_*Ca*_) of osteocytes responsive to per unit load, multiplied by the number of calcium spikes (*n*_*Ca*_) per unit load) is approximately, directly proportional to the dissipation energy density, *ϕ*. Thus dissipation energy density has a biological basis, as it can be directly related to the calcium oscillations crucial for new bone formation. Note, however, that the obtained *τ*_4_ = 0.2145 s here in this section is different from that (0.0202 s) obtained in Section 4.1. This may be due to difference in bone poroelastic / viscoelastic properties due to different mouse strains (C57BL/6 mice [35] vs. genetically engineered OtGP3 mice [38]) and different bones (axially loaded ulna [35] vs. three-point bending of 3rd metatarsal [38]). In addition, the work by Lewis et al. [38] used surgical procedure to expose the third metatarsal bone, which was kept inside a 37°C PBS-filled bath while being loaded and observed for calcium ion oscillation, which may also bring about the difference. The model, however, captures the trend, as *p*_*Ca*_ first increases with frequency and then decreases with the frequency, in agreement with the experimental results by Lewis et al. [38]. A detailed analysis in this respect has also been taken as the future work.

### 4.3. Application to Physiological Loading

The estimated strain in mouse tibia due to normal gait is 368 ± 30 με, as per Prasad et al. [39]. Assuming this strain to be predominant; as per Eq. (102), this strain is a threshold strain for *f* =(*C*/*ϵ*_*thres*_)^2^= 24.73 Hz of predominant loading frequency, which does correspond to a normal muscle stimulation frequency (20Hz to 30 Hz) required for high force generation [26], [40].

### 4.4. Incorporating Number of Loading Cycles

The dissipation energy density for 1 cycle of loading is *ϕ*. The total dissipation energy density corresponding to *N* cycles of loading is given by

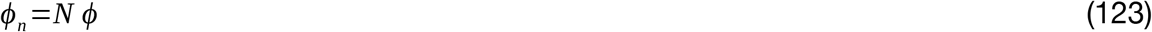

The total reference or threshold dissipation energy density corresponding to its *N* cycles is also given by

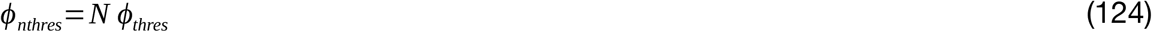

As per Eq. (99),

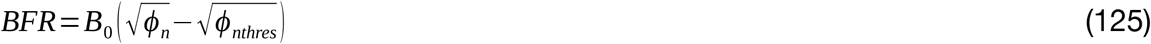

where *B*_0_ is the constant *B* for only one loading cycle.

From Eqs. (123) and (124),

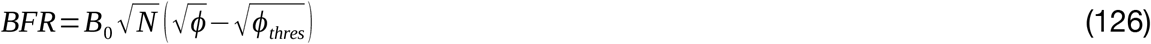

or

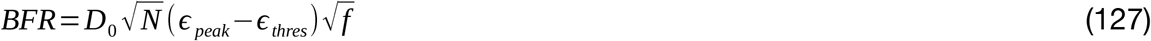

where *D*_0_ is the constant *D* for only one cycle of loading.

The value of *D* computed in Section 3.3 i.e. *D* = 0.0508 μm-s^1/2^/με/year, is for 360 cycles per day [7]; thus

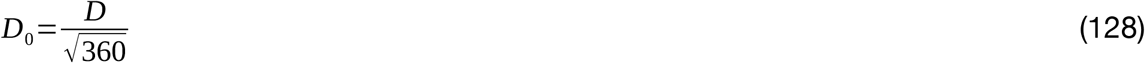

or *D*_0_ = 2.677×10^−3^ μm-s^1/2^/με/year

As per Eq. (127), if the loading waveform is kept the same except for the number of the cycles, then:

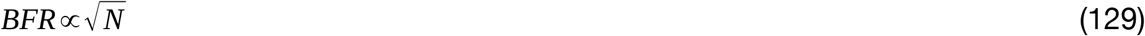

or

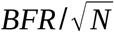 is a constant, in that case.

Srinivasan et al. [11] used a trapezoidal loading waveform. For peak strain 1250 με, the BFR values were 0.042 ± 0.026, 0.100 ± 0.032, 0.154 ± 0.054 μm/day, respectively, for 10, 50 and 250 cycles per day. The corresponding 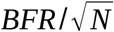 values are 0.0133 ± 0.0082, 0.0141 ± 0.0045, 0.0097 ± 0.0034 μm/day, respectively, which are not significantly different from the mean value of 0.01238 μm/day (p = 0.954, Chi Square Goodness of Fit test). Thus 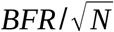 is indeed found to be a constant for a given loading protocol.

### 4.5. Incorporating Non-Sinusoidal Waveform

Non-sinusoidal waveforms such as trapezoidal loading and rest-inserted loading have been extensively used in experimental studies [11]. Dissipation energy density can be analytically computed by expressing the loading waveform as a Fourier series, as done by Prasad and Goyal [12]. The stress, strain and dissipation energy density can then be computed using their relations, as described in Section 2. Alternatively, dissipation energy density may be computed by finite element analysis, as done by Singh et al. [33], who have predicted BFR based on dissipation energy density itself.

### 4.6. Prediction of Site-Specific Adaptation on a Bone Section

The mineral apposition rate (MAR) may be formulated similar to BFR as a function of dissipation energy density, as done by Singh et al. [33], who have derived, validated and predicted the site-specific MAR based on dissipation energy density.

### 4.7. Extension to Disuse-Induced Bone Loss

As per Robinson et al. [41], a mouse walks about 377 ± 41 m every day (inside home cage). Wooley et al. [42] reported that a 2-month old mouse has stride length of 55.6 ± 1.0 mm. Thus the mouse takes about 6781 steps in a day. Due to transient muscle paralysis, dissipation energy density *ϕ* = 0. Hence, from Eqs. (99) and (102)-(104),

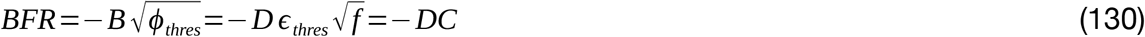

where *D* = 0.0508 μm-s^1/2^/με/year (Section 3.3) is for 360 cycles per day [7]; *C* = 1830 με/s^1/2^.

BFR/BS for *N* = 6781 cycles per day is given by: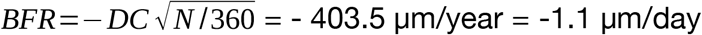, where the negative sign represents bone resorption. The value is not significantly different from the experimental value of the Bone Resorption Rate (per unit bone surface) of 1.15 ±0.67 μm/day (p = 0.95, t-test) for the distal section considered by Ausk et al. [43], [44].

## 5. Conclusions

This study derives peak strain, strain threshold and bone formation rate (BFR) from basic principles for the first time and successfully validates against the existing experimental data. The work theoretically establishes for the first time that (i) Dissipation energy density is maintained as a universal constant value in bone, (ii) If the dissipation energy density is disturbed by application of increased forces on bone, the bone acts to bring the dissipation energy density to the universal value, (iii) The corresponding bone formation rate (BFR) is proportional to increase in the square root of dissipation energy density from corresponding universal value, (iv) BFR slowly decreases to zero as dissipation energy density slowly approaches the universal value, (v) The model has a strong biological basis, and (vi) It is versatile enough to fit many of the existing works in the literature, and to be applicable to not only new bone formation, but also disuse-based bone resorption.

## Declarations

### Conflicts of interest/Competing interests

None

### Availability of data and material

The data are taken from the previously published journal articles and hence publicly available.

## Notes

### Competing Interest Statement

The authors have declared no competing interest.

### Summary of Updates

A more robust viscoelatic model has been incorporated and explained using two new figures added for better clarity. More rigorous analysis has been carried out using additional equations. The original results are now more convincing.

## References

[1] J. Wolff, “Ueber die innere Architectur der Knochen und ihre Bedeutung für die Frage vom Knochenwachsthum,” Arch. Für Pathol. Anat. Physiol. Für Klin. Med., vol. 50, pp. 389–450, 1870.

[2] H. M. Frost, “Bone ‘mass’ and the ‘mechanostat’: a proposal,” Anat. Rec., vol. 219, no. 1, pp. 1–9, Sep. 1987, doi: 10.1002/ar.1092190104.

[3] C. T. Rubin and L. E. Lanyon, “Regulation of bone mass by mechanical strain magnitude,” Calcif. Tissue Int., vol. 37, no. 4, pp. 411–417, Jul. 1985, doi: 10.1007/BF02553711.

[4] S. Srinivasan, B. J. Ausk, S. L. Poliachik, S. E. Warner, T. S. Richardson, and T. S. Gross, “Rest-inserted loading rapidly amplifies the response of bone to small increases in strain and load cycles,” J. Appl. Physiol. Bethesda Md 1985, vol. 102, no. 5, pp. 1945–1952, May 2007, doi: 10.1152/japplphysiol.00507.2006.

[5] S. Srinivasan, D. A. Weimer, S. C. Agans, S. D. Bain, and T. S. Gross, “Low-magnitude mechanical loading becomes osteogenic when rest is inserted between each load cycle,” J. Bone Miner. Res. Off. J. Am. Soc. Bone Miner. Res., vol. 17, no. 9, pp. 1613–1620, Sep. 2002, doi: 10.1359/jbmr.2002.17.9.1613.

[6] C. H. Turner, M. R. Forwood, J. Y. Rho, and T. Yoshikawa, “Mechanical loading thresholds for lamellar and woven bone formation,” J. Bone Miner. Res. Off. J. Am. Soc. Bone Miner. Res., vol. 9, no. 1, pp. 87–97, Jan. 1994, doi: 10.1002/jbmr.5650090113.

[7] Y. F. Hsieh and C. H. Turner, “Effects of loading frequency on mechanically induced bone formation,” J. Bone Miner. Res. Off. J. Am. Soc. Bone Miner. Res., vol. 16, no. 5, pp. 918–924, May 2001, doi: 10.1359/jbmr.2001.16.5.918.

[8] Y. F. Hsieh, A. G. Robling, W. T. Ambrosius, D. B. Burr, and C. H. Turner, “Mechanical loading of diaphyseal bone in vivo: the strain threshold for an osteogenic response varies with location,” J. Bone Miner. Res. Off. J. Am. Soc. Bone Miner. Res., vol. 16, no. 12, pp. 2291– 2297, Dec. 2001, doi: 10.1359/jbmr.2001.16.12.2291.

[9] T. Sugiyama, L. B. Meakin, W. J. Browne, G. L. Galea, J. S. Price, and L. E. Lanyon, “Bones’ adaptive response to mechanical loading is essentially linear between the low strains associated with disuse and the high strains associated with the lamellar/woven bone transition,” J. Bone Miner. Res. Off. J. Am. Soc. Bone Miner. Res., vol. 27, no. 8, pp. 1784– 1793, Aug. 2012, doi: 10.1002/jbmr.1599.

[10] S. Srinivasan et al., “Static Preload Inhibits Loading-Induced Bone Formation,” JBMR Plus, vol. 3, no. 5, p. e10087, May 2019, doi: 10.1002/jbm4.10087.

[11] S. Srinivasan et al., “Rescuing Loading Induced Bone Formation at Senescence,” PLoS Comput. Biol., vol. 6, no. 9, p. e1000924, Sep. 2010, doi: 10.1371/journal.pcbi.1000924.

[12] J. Prasad and A. Goyal, “An Invertible Mathematical Model of Cortical Bone’s Adaptation to Mechanical Loading,” Sci. Rep., vol. 9, no. 1, p. 5890, Apr. 2019, doi: 10.1038/s41598-019-42378-5.

[13] D. R. Carter, D. P. Fyhrie, and R. T. Whalen, “Trabecular bone density and loading history: regulation of connective tissue biology by mechanical energy,” J. Biomech., vol. 20, no. 8, pp. 785–794, 1987, doi: 10.1016/0021-9290(87)90058-3.

[14] A. G. Robling and C. H. Turner, “Mechanical signaling for bone modeling and remodeling,” Crit. Rev. Eukaryot. Gene Expr., vol. 19, no. 4, pp. 319–338, 2009, doi: 10.1615/critreveukargeneexpr.v19.i4.50.

[15] A. K. Tiwari and J. Prasad, “Computer modelling of bone’s adaptation: the role of normal strain, shear strain and fluid flow,” Biomech. Model. Mechanobiol., vol. 16, no. 2, pp. 395– 410, Apr. 2017, doi: 10.1007/s10237-016-0824-z.

[16] C. Kumar, I. Jasiuk, and J. Dantzig, “Dissipation energy as a stimulus for cortical bone adaptation,” J. Mech. Mater. Struct., vol. 6, no. 1–4, pp. 303–319, Jun. 2011, doi: 10.2140/jomms.2011.6.303.

[17] C. H. Turner, “Three rules for bone adaptation to mechanical stimuli,” Bone, vol. 23, no. 5, pp. 399–407, Nov. 1998, doi: 10.1016/s8756-3282(98)00118-5.

[18] A. K. Tiwari, A. Goyal, and J. Prasad, “Modeling cortical bone adaptation using strain gradients,” Proc. Inst. Mech. Eng. [H], vol. 235, no. 6, pp. 636–654, Jun. 2021, doi: 10.1177/09544119211000228.

[19] Y.-F. Hsieh, T. Wang, and C. H. Turner, “Viscoelastic response of the rat loading model: implications for studies of strain-adaptive bone formation,” Bone, vol. 25, no. 3, pp. 379–382, Sep. 1999, doi: 10.1016/S8756-3282(99)00181-7.

[20] M. A. Biot, “General Theory of Three-Dimensional Consolidation,” J. Appl. Phys., vol. 12, no. 2, pp. 155–164, Feb. 1941, doi: 10.1063/1.1712886.

[21] W. Thomson, “IV. On the elasticity and viscosity of metals,” Proc. R. Soc. Lond., vol. 14, pp. 289–297, Dec. 1865, doi: 10.1098/rspl.1865.0052.

[22] W. Voigt, “Ueber die innere Reibung der festen Körper, insbesondere der Krystalle,” Abh. K. Ges. Von Wiss. Zu Gött., vol. 36, pp. 3–47, 1890.

[23] F. P. Beer, E. R. Johnston, J. T. DeWolf, D. F. Mazurek, and S. Sanghi, Mechanics of materials, Seventh edition in SI Units, Special India edition. New Delhi: McGraw-Hill Education (India) Private Limited, 2017.

[24] S. A. El Sayed, T. A. Nezwek, and M. Varacallo, “Physiology, Bone,” in StatPearls, Treasure Island (FL): StatPearls Publishing, 2024. Accessed: Aug. 21, 2024. [Online]. Available: http://www.ncbi.nlm.nih.gov/books/NBK441968/

[25] E. Kreyszig, H. Kreyszig, and E. J. Norminton, Advanced engineering mathematics, 10th ed. Hoboken, NJ: John Wiley, 2011.

[26] C. R. Ethier and C. A. Simmons, Introductory biomechanics: from cells to organisms, 13th printing. in Cambridge texts in biomedical engineering. Cambridge: Cambridge University Press, 2018.

[27] R. S. Lakes, J. L. Katz, and S. S. Sternstein, “Viscoelastic properties of wet cortical bone—I. Torsional and biaxial studies,” J. Biomech., vol. 12, no. 9, pp. 657–678, Jan. 1979, doi: 10.1016/0021-9290(79)90016-2.

[28] K. Levenberg, “A method for the solution of certain non-linear problems in least squares,” Q. Appl. Math., vol. 2, no. 2, pp. 164–168, 1944, doi: 10.1090/qam/10666.

[29] D. W. Marquardt, “An Algorithm for Least-Squares Estimation of Nonlinear Parameters,” J. Soc. Ind. Appl. Math., vol. 11, no. 2, pp. 431–441, Jun. 1963, doi: 10.1137/0111030.

[30] K. Pearson, “On the criterion that a given system of deviations from the probable in the case of a correlated system of variables is such that it can be reasonably supposed to have arisen from random sampling,” Lond. Edinb. Dublin Philos. Mag. J. Sci., vol. 50, no. 302, pp. 157– 175, Jul. 1900, doi: 10.1080/14786440009463897.

[31] R. Clausius, “Ueber die bewegende Kraft der Wärme und die Gesetze, welche sich daraus für die Wärmelehre selbst ableiten lassen,” Ann. Phys. Chem., vol. 155, no. 4, pp. 500–524, 1850, doi: 10.1002/andp.18501550403.

[32] J. R. Cho, H. W. Lee, W. B. Jeong, K. M. Jeong, and K. W. Kim, “Numerical estimation of rolling resistance and temperature distribution of 3-D periodic patterned tire,” Int. J. Solids Struct., vol. 50, no. 1, pp. 86–96, Jan. 2013, doi: 10.1016/j.ijsolstr.2012.09.004.

[33] S. Singh, S. J. Singh, and J. Prasad, “Derivation, validation, and prediction of loading-induced mineral apposition rates at endocortical and periosteal bone surfaces based on fluid velocity and pore pressure,” Bone Rep., vol. 19, p. 101729, Dec. 2023, doi: 10.1016/j.bonr.2023.101729.

[34] Student, “The Probable Error of a Mean,” Biometrika, vol. 6, no. 1, p. 1, Mar. 1908, doi: 10.2307/2331554.

[35] S. J. Warden and C. H. Turner, “Mechanotransduction in the cortical bone is most efficient at loading frequencies of 5–10 Hz,” Bone, vol. 34, no. 2, pp. 261–270, Feb. 2004, doi: 10.1016/j.bone.2003.11.011.

[36] K. J. Lewis, “Osteocyte calcium signaling – A potential translator of mechanical load to mechanobiology,” Bone, vol. 153, p. 116136, Dec. 2021, doi: 10.1016/j.bone.2021.116136.

[37] D. Jing et al., “In situ intracellular calcium oscillations in osteocytes in intact mouse long bones under dynamic mechanical loading,” FASEB J., vol. 28, no. 4, pp. 1582–1592, Apr. 2014, doi: 10.1096/fj.13-237578.

[38] K. J. Lewis et al., “Osteocyte calcium signals encode strain magnitude and loading frequency in vivo,” Proc. Natl. Acad. Sci., vol. 114, no. 44, pp. 11775–11780, Oct. 2017, doi: 10.1073/pnas.1707863114.

[39] J. Prasad, B. P. Wiater, S. E. Nork, S. D. Bain, and T. S. Gross, “Characterizing gait induced normal strains in a murine tibia cortical bone defect model,” J. Biomech., vol. 43, no. 14, pp. 2765–2770, Oct. 2010, doi: 10.1016/j.jbiomech.2010.06.030.

[40] C. T. Rubin, K. J. McLeod, and S. D. Bain, “Functional strains and cortical bone adaptation: Epigenetic assurance of skeletal integrity,” J. Biomech., vol. 23, pp. 43–54, Jan. 1990, doi: 10.1016/0021-9290(90)90040-A.

[41] L. Robinson, A. Plano, S. Cobb, and G. Riedel, “Long-term home cage activity scans reveal lowered exploratory behaviour in symptomatic female Rett mice,” Behav. Brain Res., vol. 250, pp. 148–156, Aug. 2013, doi: 10.1016/j.bbr.2013.04.041.

[42] C. M. Wooley, S. Xing, R. W. Burgess, G. A. Cox, and K. L. Seburn, “Age, experience and genetic background influence treadmill walking in mice,” Physiol. Behav., vol. 96, no. 2, pp. 350–361, Feb. 2009, doi: 10.1016/j.physbeh.2008.10.020.

[43] B. J. Ausk, P. Huber, S. L. Poliachik, S. D. Bain, S. Srinivasan, and T. S. Gross, “Cortical bone resorption following muscle paralysis is spatially heterogeneous,” Bone, vol. 50, no. 1, pp. 14– 22, Jan. 2012, doi: 10.1016/j.bone.2011.08.028.

[44] H. Shekhar, S. Singh, and J. Prasad, “Predicting Cortical Bone Resorption in Disuse Condition Caused by Transient Muscle Paralysis,” Nov. 21, 2023. doi: 10.1101/2023.11.21.568078.

